# A pipeline for validation of sRNA effectors suggests cross-kingdom communication in the symbiosis of *Arabidopsis* with *Serendipita indica*

**DOI:** 10.1101/2023.12.19.572297

**Authors:** Sabrine Nasfi, Saba Shahbazi, Katharina Bitterlich, Ena Šečić, Karl-Heinz Kogel, Jens Steinbrenner

## Abstract

Bidirectional communication between pathogenic microbes and their plant hosts via small (s)RNA-mediated cross-kingdom RNA interference (ckRNAi) is one key element for successful host colonisation. However, whether mutualistic fungi from the Serendipitaceae family, known for their extremely broad host range, employ sRNAs to colonize plant roots is still under discussion. To address this question, we developed a pipeline to validate the accumulation, translocation, and activity of fungal sRNAs in post-transcriptional silencing of *Arabidopsis thaliana* genes. Using stem-loop PCR, we detected the expression of a specific set of *Serendipita indica* (*Si)*sRNAs, targeting host genes involved in cell wall organization, hormonal signalling regulation, immunity, and gene regulation. To confirm the gene silencing activity of these sRNA in plant cells, *Si*sRNAs were transiently expressed in protoplasts. Stem-loop PCR proved the expression of sRNAs, while qPCR validated post-transcriptional gene silencing of their predicted target genes. Furthermore, ARGONAUTE 1 immunoprecipitation (*At*AGO1-IP) revealed the loading of fungal *Si*sRNAs into the plant RNAi machinery, suggesting the translocation of *Si*sRNA from the fungus into root cells. In conclusion, this study provides a blueprint for rapid selection and analysis of sRNA effectors in plant-microbe interactions and further suggests cross-kingdom communication in the Sebacinoid symbiosis.

**Highlight:** Small RNAs of the beneficial fungus *Serendipita indica* are translocated and silence Arabidopsis genes at the onset of the interaction, revealing cross-kingdom communication in sebacinoid symbiosis.

## Introduction

RNA interference (RNAi) is a regulatory process in most eukaryotes in which gene expression is silenced at the transcriptional or post-transcriptional level by the action of small (s) RNA; it is associated with the control of genome stability, developmental processes, and responses to biotic and abiotic stresses (Zhan and Meyers, 2023). Two types of sRNAs predominate in plants: micro (mi) RNAs are 20-22 nucleotides (nt) in length, processed from primary miRNAs (pri-miRNAs) containing a stem-loop structure encoded by microRNA (*MIR)* genes (Meyers and Axtell, 2019; Reinhart et al., 2002). In contrast, small-interfering (si) RNAs are RNA molecules of 20-24 nt in length produced from longer endogenous RNA templates or from exogenous RNA sequences (viruses or transgenes). miRNA and siRNA precursors are processed by RNase III-like endonucleases named DICER-like (DCL), then bound by ARGONAUTE (AGO) proteins, which are incorporated into the RNA-Induced Silencing Complex (RISC), where they target complementary mRNA transcripts to induce post-transcriptional gene silencing (PTGS) (Iwakawa & Tomari, 2022). Plant sRNAs can also move intercellularly across plasmodesmata and systemically across the phloem, acting cell non-autonomously and can also translocate between interacting species (Zhan & Meyers, 2023). In the latter case, sRNAs are transferred bidirectionally between plant hosts and pathogenic microbes to modulate defence and virulence, a phenomenon called cross-kingdom RNAi (ckRNAi) (Göhre and Weiberg, 2023; Mahanty et al., 2023; Wang et al., 2017). ckRNAi in plant-pathogen interactions was discovered in 2013 when the necrotrophic fungus *Botrytis cinerea* was shown to induce silencing of host defence genes in *Solanum lycopersicum* and *Arabidopsis thaliana* (*At*) (Weiberg et al., 2013). Since then, several reports supported the existence of ckRNAi (Wang & Dean, 2020) and showed that sRNA effectors exchange can be bidirectional (Zhang et al., 2016; Cai et al., 2018).

Already in 2006, Huang and colleagues reported reduced nematode infectivity when root-knot nematodes (RKN) fed on roots of Arabidopsis plants expressing double-stranded (ds) RNA targeting the *16D10* gene, which encodes a conserved secreted RKN root growth-stimulating peptide known to be involved in parasitism (Huang et al., 2006). Later, it was shown that sRNA can indeed move from a plant to an attacking pathogen in the interaction of barley and wheat with the powdery mildew fungus (*Blumeria graminis*) (Nowara et al., 2010). Transient transformation with a plasmid overexpressing dsRNA directed against the fungal *AVR10* gene increased the susceptibility of leaf epidermal cells to the corresponding *Blumeria graminis* races, a technique termed host-induced-gene-silencing (HIGS). Since then, numerous reports have shown that HIGS is an efficient strategy for pests and disease control in crops (Cai et al., 2018; Koch et al., 2013; Liu et al., 2020; Sang & Kim, 2020; Zand Karimi & Innes, 2022).

Although ckRNAi and related artificial processes such as HIGS were discovered in plants more than 10 years ago, the molecular mechanisms are still not fully understood. For instance, the mode of bidirectional transfer of RNAs in extracellular vesicles (EVs) is still a matter of debate (He, Cai, et al., 2021; He, Hamby, et al., 2021; Nasfi & Kogel, 2022; Rutter & Innes, 2018), and the occurrence in mutualistic symbioses is little researched albeit a growing number of studies are shedding light on ckRNAi in plant-symbiont interactions (Qiao et al., 2023).

ckRNAi in mycorrhizal symbioses seems to include miRNA. For example, Pmic_miR-8, a miRNA from the ectomycorrhizal fungus *Pisolithus microcarpus* enhanced the colonization of its host *Eucalyptus grandis and* thus plays a role in the beneficial ectomycorrhizal symbiosis (Wong-Bajracharya et al., 2022). In addition, recent *in silico* analysis of the sRNA population from *Rhizophagus irregularis*, supported by degradome data from *Medicago truncatula*, has shown that some sRNAs of Rhizophagus modulate root colonization (Silvestri et al., 2019). Moreover, tRFs identified in root nodule symbioses between soybean and the bacterial symbiont *Bradyrhizobium japonicum* hijack *Glycine max* AGO1 and are positive regulators of rhizobial infection and nodule formation (Ren et al., 2019), further emphasising the existence of ckRNAi in plant-mutual interactions.

Key to understanding ckRNAi is the identification and functional validation of translocated sRNA (sRNA effectors), and tools are needed to study the mode of action of numerous sRNAs derived from extensive sRNA sequencing data. Here, we report on a pipeline for sRNA effectors validation by studying the interaction between the mutualistic basidiomycete fungus *Serendipita indica* (*Si*) and the dicot model plant *Arabidopsis thaliana* (*At*). The fungus was originally isolated from the roots of the shrubs *Prosopis juliflora* and *Ziziphus nummularia* in the Indian Thar desert (Verma et al., 1998). Its beneficial activity includes plant growth promotion, plant resistance priming, enhanced nitrate and phosphate delivery, promotion of adventitious root and root hair formation, early flowering, support of higher seed yield, alteration in secondary metabolites, and hardening of tissue cultured plants (Glaeser et al., 2016; Qiang et al., 2012; N. Verma et al., 2022; Weiß et al., 2016; Xu et al., 2018; Zuccaro et al., 2011). The fungus transfers protein effectors to host cells to exploit the host’s metabolism and promote microbial colonization (Akum et al., 2015; Osborne et al., 2023). To further explore the molecular basis of the mutualistic interaction formed by *Si* with a wide range of plants (Sebacinalean symbiosis), we recently demonstrated the global change in sRNA profiles in the interaction of the fungus and the grass model *Brachypodium distachyon* (*Bd*) (Šečić et al., 2021). Among *Bd*- and *Si*-generated sRNAs with putative functions in the interacting organism, we found proteins involved in cell wall organization, hormonal signalling regulation and immunity as potential targets of putative fungal sRNA effectors.

Building upon the findings from our prior work (Šečić et al., 2021), we have selected a set of fungal sRNAs with interesting predicted *At* targets to further investigate their activity in the *Si-At* interaction. We developed a functional protoplast assay to validate potential putative fungal sRNA effectors assessing their gene-silencing activity. Using stem-loop PCR, we confirmed sRNA transformation and expression in plant protoplasts and recorded the downregulation of predicted host target genes via qPCR. Additionally, we investigated the capability of *Si*sRNA effector candidates in mediating the degradation of host mRNA by 5’-RLM-RACE. Finally, *At*AGO1 immunoprecipitation assay, confirmed the loading of fungal *Si*sRNAs into the plant’s RNAi machinery. The data support the existence of naturally occurring ckRNAi between the mutualistic fungus *Serendipita indica* and *Arabidopsis thaliana*.

## Materials and methods

### Plant material and growth conditions

For protoplast isolation, *Arabidopsis thaliana* plants of ecotype Columbia-0 (Col-0) were cultivated in a 4:1 ratio of T-type soil (F.-E. Typ Nullerde, Hawita) and crystal quartz sand mixture for 4 to 5 weeks in a growth chamber in a 19°C day/18°C night (9 h of light, 150 μmol photons m^-2^ s^-1^) and a constant relative humidity of 60%. For interaction studies of *At* roots and *Serendipita indica* (IPAZ-11827, Institute of Phytopathology, Giessen, Germany), plants were grown on vertical square Petri dishes on an ATS medium (Lincoln et al., 1990) without sucrose and supplemented with 4.5 g/L Gelrite (Duchefa #G1101) in a 22°C day/18°C night cycle (8 h of light). Roots of 14-day-old plants were inoculated with 1 mL of a suspension of 500,000 chlamydospores/mL in aqueous 0.02% Tween 20 per Petri dish.

### sRNA sequencing and Arabidopsis target prediction

In our previous study (Šečić et al., 2021) we investigated changes in mRNA and sRNA expression profiles during the mutualistic interaction between *Serendipita indica* and the grass model *Brachypodium distachyon* using high throughput sRNAseq and transcriptome analysis. Raw sRNA sequencing data underwent bioinformatic filtering steps, as detailed in Šečić *et al* 2021, and were aligned to the genomes and transcriptomes of both organisms. *Si*sRNAs were categorized into two groups: endogenous *Si*sRNAs present in axenic fungal culture, and putative ck-*Si*sRNAs exhibiting higher abundance in the colonized *Bd* roots, therefore potentially playing a crucial role in the establishment of RNAi mediated mutualism. Size distribution analysis of the putative ck-*Si*sRNAs revealed a prominent peak of 21 nt molecules in the *Si-Bd* samples. By focusing on 21 nt *Si*sRNAs, we identified 412 unique induced putative ck-sRNAs, of which carry 30% adenine (A), 26% uracil (U), 21% cytosine (C) and 22% guanine (G) at their 5’-end.

### Plasmid construct

*At*MIR390a distal stem-loop was digested from the pMDC32B-*At*MIR390a-B/c vector (Plasmid #51776, https://www.addgene.org, Carbonell et al., 2014) using Fast digest enzymes EcoRI and HindIII (Thermo Fisher Scientific, FD0274 & FD0505). The pUC18-*At*MIR390a-B/c construct was generated by ligating the *At*MIR390a precursor into the pUC18 vector (plasmid #50004) backbone using T4 DNA Ligase (Thermo Fischer Scientific, EL0011). The *RFP* insert was digested from the pBeaconRFP_GR vector (https://gatewayvectors.vib.be/index.php/ ID: 3_20, Bargmann & Birnbaum, 2009) using the restriction enzyme NdeI (NEB, R0111S) and further ligated into the NdeI site of the pUC18-*At*MIR390a-B/c vector. 75mer oligonucleotides comprising the amiRNAs and *Si*sRNAs sequences were cloned into the BsaI-sites by Golden Gate strategy using BsaI-HF^®^v2 (NEB, R3733) and T4 DNA Ligase (Thermo Fischer Scientific, EL0011) in a single reaction, replacing the toxic ccdB gene which allows the selection of 75-mer-positive clones. Before cloning 75-mer oligonucleotides, a third BsaI-site in the pUC18 backbone which would interfere with the Golden-Gate cloning was removed by site-directed mutagenesis using Phusion High-Fidelity DNA polymerase (Thermo Fisher Scientific, F553S) and the Fast digest Dpn I endonuclease (Thermo Fisher Scientific, FD1703) as well as custom-designed site-directed mutagenesis primers pUC18-Mut-Fwd and pUC18-Mut-Rev.

The artificially designed amiRNAs, their 5’-nucleotide, their predicted targets, their target aligned fragment, and their expectation value are listed in Supplemental Tables 1. Same details of the selected putative ck-*Si*sRNAs cloned in the expression vector for further PTGS analysis are listed in Supplemental Table 2. Supplemental Figure 1 illustrate an alignment between amiRNAs and *Si*sRNA and their predicted selected targets as displayed by psRNATarget web-tool. 75-mer oligonucleotides for amiRNAs and *Si*sRNAs were designed following the P-SAMS algorithm developed by the James Carrington lab (Fahlgren et al., 2016) available at http://p-sams.carringtonlab.org. 75 mer oligonucleotides of amiRNAs and *Si*sRNAs are listed in Supplemental Table 3. An *in-silico* cloning design was performed using Snapgene software (www.snapgene.com) and the final pUC18-*At*MIR390a-RFP-sRNA vector map is shown in Supplement Figure 2.

### Protoplast transformation

The “Tape-Arabidopsis Sandwich” method was used for protoplast isolation (Wu et al., 2009). The transformation was performed following the transient expression of recombinant genes using the Arabidopsis mesophyll protoplast (TEAMP) approach (Yoo et al., 2007) with minor modifications. Since the transformation of protoplasts was key to all subsequent experiments, we attempted to optimize its efficiency by varying *i.* protoplast concentration (10x 10^4^ and 5x 10^4^ protoplasts/mL), *ii.* plasmid concentration (20 µg, 30 µg, and 40 µg) pUC18-*At*MIR390a-RFP-sRNA and *iii*. incubation time (24 and 48 h). High transformation efficiency was reported when using 5x 10^4^ protoplasts/ml with a plasmid concentration of 30 µg and an incubation time of 24 h, which was comparable to the reported transformation efficiencies of *At* leaf protoplasts (Yoo et al., 2007). Total cell numbers were counted using a Fuchs-Rosenthal counting chamber under an optical microscope. A microscopic check of protoplast transformation was done using an epifluorescence microscope (TCS SP2 Leica). Images were acquired with the image software Leica Application Suite (LAS) and all counting was performed in triplicates. Transformed protoplasts were stored at −80°C for further studies.

### RNA extraction and quality analysis

Total RNA was extracted from protoplasts using QuickRNA™ Miniprep Kit (Zymo Research, R1050) with an on-column DNase I treatment. Total RNA from *Si-At* interaction growing on ATS plates was extracted using Direct-zol RNA Miniprep (Zymo Research, R2051) with an on-column DNase I treatment. The RNA concentration was determined with the NanoDrop ND-1000 Spectrophotometer (Thermo Fisher Scientific, USA) and the purity was determined by measuring A260/280 and A260/230 ratios. The quality of the RNA extracted from transformed protoplasts was checked using the Agilent 2100 Bioanalyzer Nano Chip (Agilent, Germany).

### Stem-Loop PCR

Designed amiRNAs and *Si*sRNAs sequences were used as templates to design specific stem-loop (SL) primers matching the corresponding sRNA over six nt at the 3’end. Hairpin primers (cDNAhp) and forward primers were designed using the tool published in Adhikari et al (2013) based on Varkonyi-Gasic et al., (2007) (Adhikari et al., 2013; Varkonyi-Gasic & Hellens, 2011). Stem-loop PCR was performed as described (Werner et al., 2021). End-point PCR was used to assess the expression of amiRNAs and *Si*sRNAs. The PCR program was optimized to 95 °C for 5 min; followed by 40 cycles of 95 C for 30 s, 60°C for 30 s, 72°C for 30 s; and 72°C for 5 min and then held at 4°C. PCR products were separated and visualized in 2% TBE-agarose gels. Stem-loop PCR primers are listed in Supplemental Table 4.

### PTGS detection by qPCR

The standard curve method was used to test the efficiency of the qPCR transcript primers using a serial dilution of *At* cDNA library along with 3 different primer concentrations (0.4 µM, 0.2 µM and 0.1 µM for each primer) and 5 µL of SybrGreen (Sigma-Aldrich). The total volume of 10 µL and three technical replicates are considered for each reaction. Before master mix preparation, 2 µL ROX (CRX reference dye, Promega, C5411) were added to 1 mL SybrGreen as a passive reference dye that allows fluorescent normalization for qPCR data. For the preparation of cDNA first strands, 500 ng of RNA samples from transformed *At* protoplasts and control protoplasts were used. qPCR was performed using the QuantStudio 5 Real-Time PCR system (Applied Biosystems) in the 384-well plate as follows: 50°C for 2 min, 95°C for 10 min, followed by 40 cycles of 95°C for 15 s, and 60°C for 1 min, with an additional melt curve stage consisting of 15 s at 95°C, 1 min at 60°C, and 15 s at 95°C.

To evaluate the relative quantification of *At* transcripts by qPCR, the gene expression level was standardized to the control protoplasts. Fold change of *ΔΔ*ct values and standard errors were calculated for all mean values. All qPCR primers are listed in Supplemental Table 5.

### 5’-RLM-RACE

RNA ligase-mediated rapid amplification of cDNA ends (5’-RLM-RACE) was performed using FirstChoice^®^ RLM-RACE kit (Thermo Fisher Scientific) following the manufacturer protocol and omitting the dephosphorylation and decapping steps. One µg of RNA isolated from *At* protoplasts and control protoplasts were used as a template and ligated to the 5’-RACE adapter using T4 RNA Ligase [10 U/µL] (Thermo Fisher Scientific). The ligation reaction was used entirely to generate the first cDNA strand. Two rounds of nested hot-start touch-down PCR were performed using outer (first) and inner (second) 5’-RLM-RACE universal primers in combination consecutively with gene outer specific and gene inner specific primers. PCR products were evaluated in a 1.5% agarose gel and bands of the expected size were excised. Products were cleaned with the Wizard^®^ SV Gel and PCR Clean-Up System (Promega) and cloned with the pGEM^®^-T Vector Systems (Promega). For each band, six clones were selected for sequencing at LGC genomics. The oligonucleotides used are listed in the Supplemental Table 6.

### T-vector cloning and sequencing

Stem-loop PCR and 5’-RLM-RACE PCR amplification products were purified using The Wizard^®^ SV Gel and PCR Clean-Up System (Promega). Cloning of the different PCR products was performed as described in the pGEM^®^-T Vector system according to the instructoŕs instructions (Promega). Sequencing was performed at LGC Genomics (Berlin, Germany) and analysed using the Snapgene tool (GSL Biotech, available at snapgene.com). All PCR primers are presented in Supplemental Table 7.

### AGO-IP

*At*AGO1-Co-immunoprecipitation (Co-IP) was performed as described in the publication of Dunker et *al.,* 2021 with modifications. Five gram of *Si* mycelium or *At* roots inoculated or uninoculated with *Si* were ground to a fine powder using a precooled mortar and pestle. To each sample, 20 mL of immunoprecipitation extraction buffer was added, and samples were centrifuged at 3,200x g for 15 min at 4°C to remove root debris. The supernatants were filtered through double-layered Miracloth and 200 µL crude extract (CE= supernatant before antibodies) was collected and combined with 50 µL 5X protein SDS loading buffer for western blot analysis. 5 µL of anti-AGO1 polyclonal antibody (Agrisera, Catalog number: AS09527) and 200 µL of protein A agarose beads (Roche, Ref: 11719408001) were added to the remaining supernatant and incubated for 2 h at 4°C on a rotation wheel. The samples were centrifuged at 200x g for 30 sec at 4°C. From the supernatants, 200 µL were removed (SN after antibodies) and 50 µL of 5x protein SDS loading buffer was added for further western blot analysis. The remaining supernatants were discarded, and 1 mL of ice-cold IP wash buffer was added to the pelleted beads and centrifuged at 200x g for 30 sec at 4°C. The pellet beads were washed 3 times and finally resuspended in 1 mL wash buffer. Resuspended pellet beads were divided into 30% for further WB analysis supplemented with 1x protein SDS loading buffer (IP fraction) and 70% was used for sRNA recovery.

### sRNA recovery from IP fraction

IP fractions were pelleted and 300 µL IP wash buffer was added together with 150 µL RNA release buffer. Samples were incubated in a thermoshaker for 15 min at 300 rpm at 65°C. 450 µL of water-saturated phenol was added and samples were vortexed for 2 min then centrifuged at 10,000x g at room temperature (RT) for 8 min. The upper aqueous phase containing the sRNAs was transferred into a low-binding RNA tube. 450 µl of chloroform/isoamyl alcohol (24:1) (Carl Roth, Germany) was added to the RNA samples. The step was repeated twice. For RNA precipitation, 0.1x volume of 3 M sodium acetate, 2.5x volume of 96% ethanol, and 20 µg RNA grade glycogen (Thermo Fisher Scientific, R0551) were added to the RNA samples and placed overnight at −20°C. Samples were pelleted at 20,000x g for 30 min at 4°C and pellets were washed with 500 µL 80% ethanol. RNA was pelleted at 20,000x g for 20 min at 4°C, ethanol was removed, and pellets were air dried until ethanol was completely evaporated. The RNA pellets were resuspended in 8 µL DEPC-treated water and stored at −80°C.

### Western Blot analysis

For western blot analysis, a 5.5% stacking gel and a 12% resolving gel were used. For each sample, 20 µL of crude extract before antibodies (CE), supernatant after pelleting agarose beads (SN), and *At*AGO1 co-IP fraction (IP) were loaded. All samples were boiled at 95°C for 5 min before loading. Gel separation was conducted at 100 V until the SDS loading buffer reached the edge of the gel (electrophoresis takes approximately 2.5 h to 3 h). For immunodetection, a PVDF blotting membrane (Carl Roth, Germany) was used. The blot was prepared in a sandwich method in the following order: 4x Whatman paper (GE Healthcare, USA), PVDF membrane, SDS-Gel, and 4x Whatman paper. Whatman papers were soaked in Towbin buffer and the membrane was activated in methanol and washed with Towbin buffer before transfer for equilibration. Blotting was performed using BioRad TransBlot Turbo at 25 Volt and 1.0 Ampere for 30 min. The Blotting sandwich was disassembled, and the membrane was washed with 1x PBS-T (0.1% Tween) and then blocked for 1 h at RT with 10 mL 1x PBS-T (0.1% Tween) and 5% (w/v) milk powder (CarlRoth, Germany). For detection of *At*AGO1 protein, the membrane was incubated overnight at 4°C on a shaker with primary antibody solution 1 µg/µL diluted 1:4000 anti-AGO1 (Agrisera, AS09 527) in 10 mL 1% milk in 1x PBS-T. The membrane was washed 4 times for 10 min on a shaker at RT with 1x PBS-T and incubated for 2 h at RT with the secondary antibody IRDye® 800CW Goat anti-Rabbit IgG (LI-COR, 926-32211) in 10 mL 1% milk in 1x PBS-T. The membrane was washed 4 times for 10 min on a shaker at RT with 1x PBS-T. For the membrane development, chemiluminescent substrates were applied according to the manufacturer’s recommendation, and images were captured using the Bio-rad ChemiDoc MP imaging system.

### Statistical Analysis

To assess the downregulation of *At* genes via qPCR, the 2−ΔΔCt method (Livak & Schmittgen, 2001) was used for the quantification of *At* target transcript. The results are expressed as values of the fold change of the transcript expression level in comparison to the control and compared using a two-sided unpaired student *t* test. Data showed a normal distribution in the Shapiro-Wilk test or D’Agostino & Pearson test.

## Results

We developed a pipeline for the validation of sRNA gene silencing activities in the mutualistic interaction of *Serendipita indica* and *Arabidopsis thaliana*, comprising i. prediction of *Si*sRNA Arabidopsis target genes, ii. validation of the gene silencing activity of *Si*sRNAs in *At* protoplasts, iii. detection of fungal sRNAs (*Si*sRNAs) in axenic culture and *Si*-colonized roots, and iv. detection of the association of *Si*sRNAs with the *At*AGO1 protein (Figure 1).

**Figure 1.**
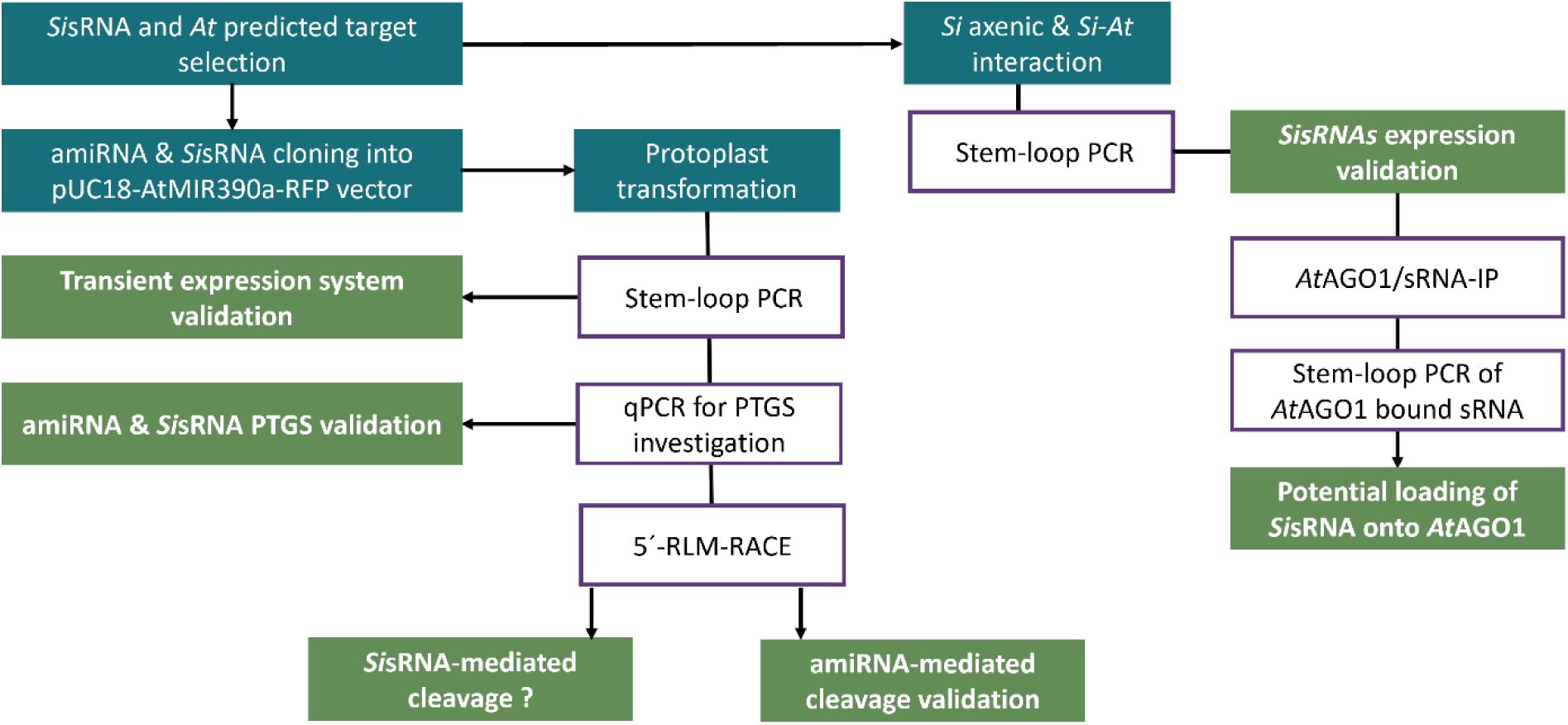
Flowchart showing the methods and outputs in this study. The turquoise colour indicates the different input material and purple indicates the wet-lab steps. Green represents the validated outputs of the study. For details also see Material and Methods. *Si*, *Serendipita indica*; *At*, *Arabidopsis thaliana*; PTGS, post-transcriptional gene silencing; amiRNA, artificial miRNA.

### *Si*sRNA selection and Prediction of Arabidopsis targets

Considering the broad host range of *Si*, we selected 14 induced ck-*Si*sRNAs from *Si* non-coding regions based on their expression levels in colonized *Bd* roots and we predicted their targets in *At* using psRNAtarget Scheme 2 (2017) default (Dai et al., 2018). Among the predicted targets, we prioritized those with intriguing descriptions and where a single *Si*sRNA has multiple target genes. Supplemental Table 8 illustrates a summary of the 14 selected putative ck-*Si*sRNA, their raw read counts normalized to *Si-*colonized *Bd* roots or to *Si* axenic, their 5’-nucleotide, and their induction level. Supplementary File 2 illustrates *At* predicted targets for the 14 selected ck-*Si*sRNAs.

### amiRNAs mediate PTGS in transformed Arabidopsis protoplasts

Arabidopsis protoplasts are a reproducible and cost-effective model system to validate the expression and the potential silencing activity of *Si*sRNA on predicted *At* target genes. To confirm the reliability of the system, we first used artificial microRNAs (amiRNAs). The use of amiRNAs allowed the amiRNA-mRNA binding to be manipulated, as they are artificially designed to have almost complete complementarity with the mRNA target gene, unlike naturally occurring *Si*sRNAs, which have varying degrees of mismatch with their predicted target genes. To this end, we designed four amiRNAs of 21 nt (amir21, amir24, amir154, and amir296) that almost fully match their predicted target genes (*AT5G25350*, *AT2G39020*, *AT4G32160,* and *AT2G45240*, respectively) and exhibit a U at their 5’-end. Subsequently, the four amiRNAs were cloned into the pUC18-*At*MIR390a-RFP vector and transformed into *At* protoplasts, reaching a transformation efficiency of 65% to 90% (calculated as the ratio between the red fluorescing protoplasts and the total number of viable protoplasts; Supplemental Figure 3). The four amiRNAs were successfully expressed in protoplast as detected by duplexed stem-loop PCR (Figure 2). Amplification products were cloned into the pGEM-T^®^ cloning vector and sequencing confirmed amiRNAs sequences.

**Figure 2.**
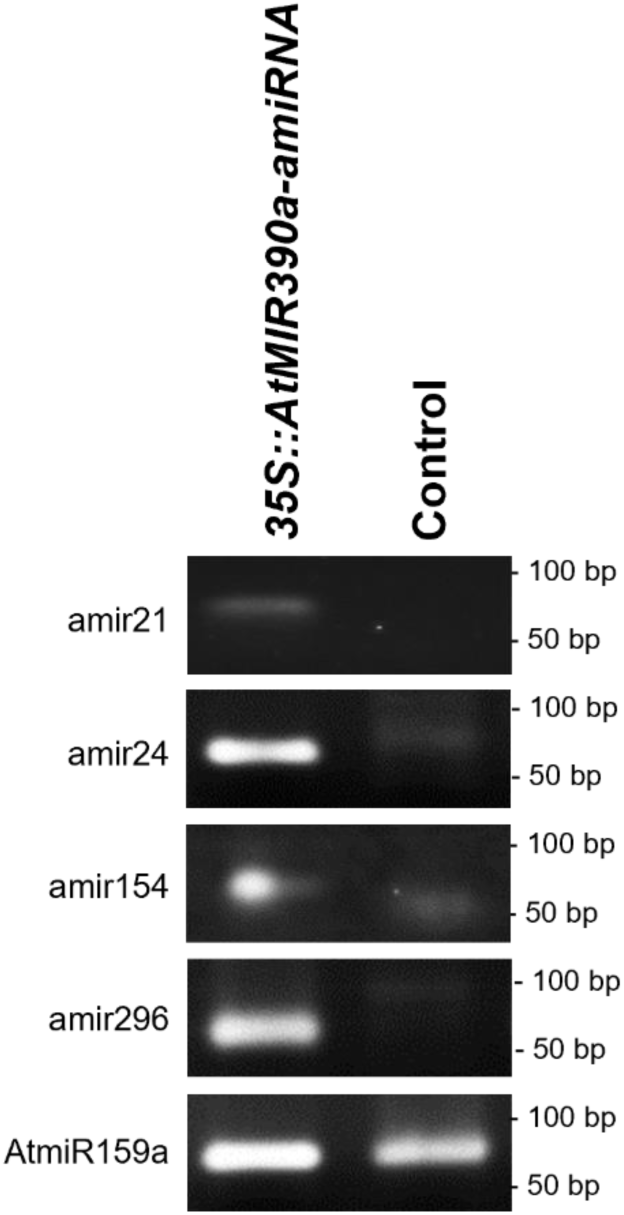
Duplexed stem-loop PCR confirming expression of amiRNAs in Arabidopsis protoplasts. Agarose gel analysis shows expression of all four amiRNAs in transformed protoplasts 24 hours post-transformation (hptr), but not in control protoplasts. Plant *At*miR159a of 21 nt length was used as a positive control.

Supplemental Table 2 provides information on these amiRNAs, including their 5’-terminal nucleotides, their predicted *At* target genes, their target aligned fragment, and their target prediction expectation values, indicating the degree of mismatches between the amiRNAs and the *At* target sequence, Supplemental Table 9 provides a description and the expression level of the predicted *At* target genes in leaf and root (Data extracted from Moreno et al., 2022).

Analysis of amir target downregulation was quantified by qPCR 24 h after the transformation of amiRNAs into protoplasts and was 68% for *AT5G25350*, 43% for *AT2G39020*, 50% for *AT4G32160*, and 65% for *AT2G4524*0, respectively (Figure 3).

**Figure 3.**
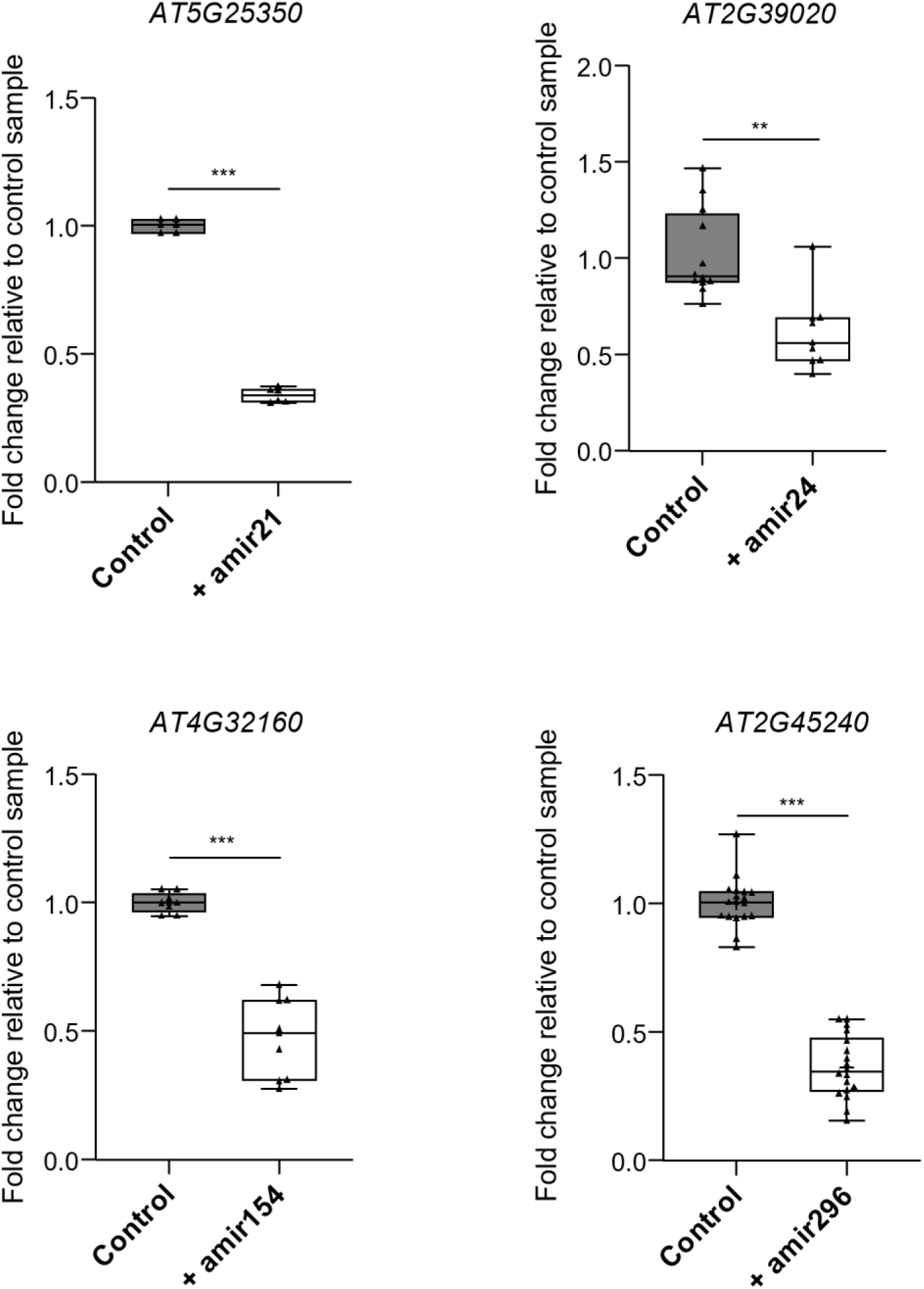
qPCR analysis of target gene silencing in Arabidopsis protoplasts 24 h post transformation with amiRNAs. The mRNA transcript levels in transformed protoplasts (+) expressing amir21, amir24, amir154, and amir296 were normalized against housekeeping gene *Ubiquitin* (*AT3G62250*) and displayed as fold change relative to control protoplasts (-) (mock-treated without amiRNA construct). Data are the average of two to three biological replicates ± standard deviation. Asterisks indicate difference at p ≤ 0.05 (*), p ≤ 0.01 (**), and p ≤ 0.001 (***) according to Student’s t-test. ns= not significant.

### 5’-RLM-RACE reveals canonical cleavage in protoplasts transformed with amiRNA

We selected amir296 and its predicted target gene *AT2G45240* to investigate the canonical PTGS cleavage site using RNA-ligase-mediated Rapid Amplification of cDNA Ends (5’-RLM-RACE) (Llave et al., 2002; Ueno et al., 2022). Samples were obtained from transformed protoplasts at 24 hptr, from control protoplasts (mock-treated without amiRNA construct) and from non-treated protoplasts. A distinct band of approximately 383 bp was amplified from protoplasts expressing amir296, consistent with the expected size of the canonical cleavage product predicted to be targeted by amir296. Unexpectedly, weaker DNA fragments of a similar size were detected in both control and non-treated protoplasts (Figure 4A). Cloning and subsequent sequencing of the amplicons confirmed that all sequences aligned with the *AT2G45240* sequence, and 50% of the obtained sequences from protoplasts transformed with pUC18-*At*MIR390a-RFP-amir296 (three of six clones) exhibited the canonical cleavage site between nucleotide 10 and 11 of amir296. In contrast, PCR products from control and non-treated protoplasts contained an *AT2G45240* fragment but did not correspond to a canonical cleavage site, suggesting a possible cleavage by an endogenous miRNA near the amir296 binding site or an RNA degradation (Figure 4B).

**Figure 4.**
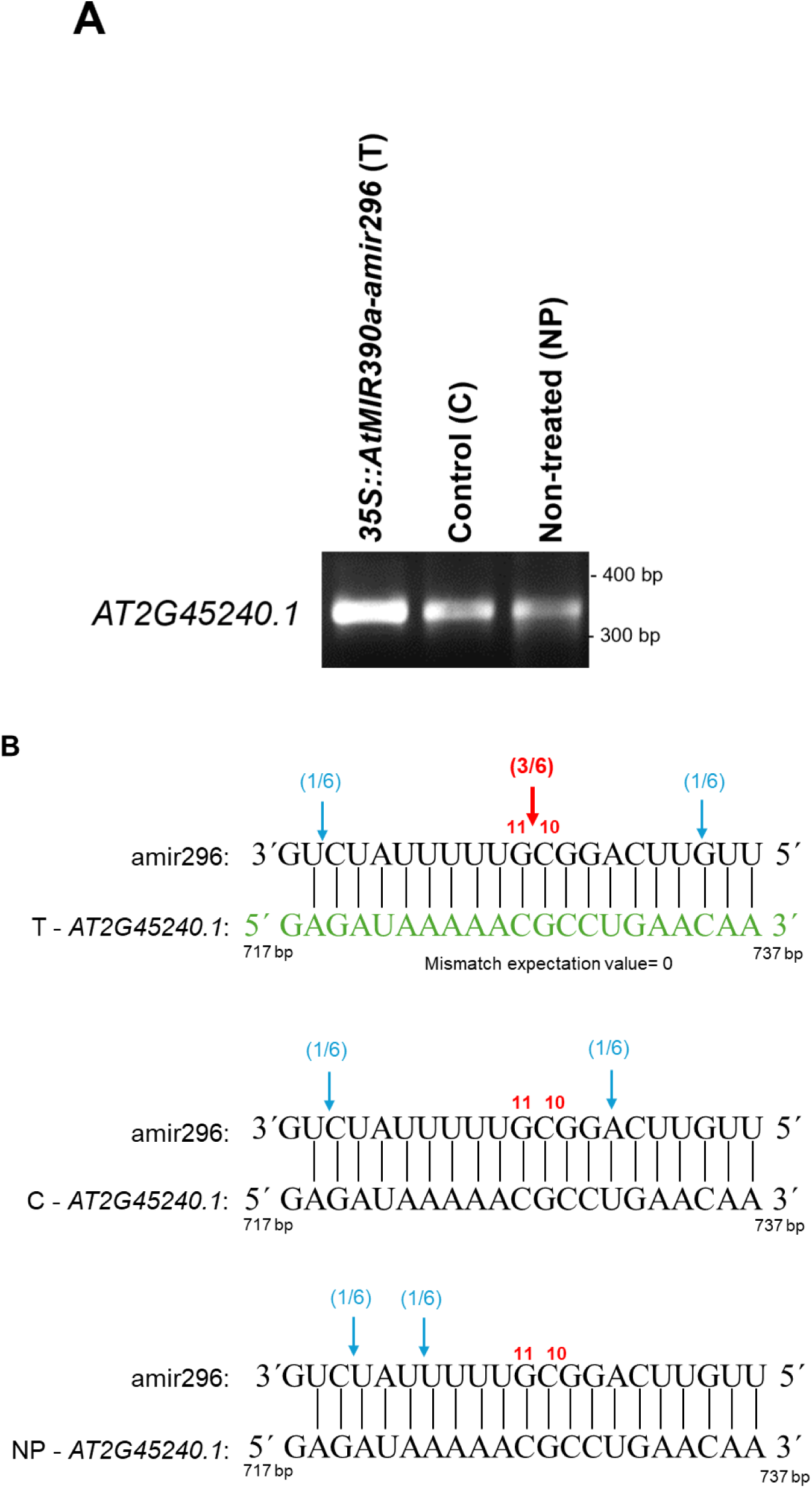
Identification of 5’-RLM-RACE target sites in *AT2G45240* mRNA following amir296 expression in Arabidopsis protoplasts (A) PCR products of the second nested 5’-RLM-RACE-PCR visualized in an EtBr-agarose gel. RNA was extracted from transformed protoplasts with amir296 (T) and from the corresponding controls (control protoplasts (C) and non-treated protoplasts (NP). (B) Mapping of *AT2G45240* target cleavage products by 5’-RLM-RACE. The predicted base pairing between amir296 and *AT2G45240* is shown in green. The red arrow indicates the detected canonical cleavage sites. The proportion of cloned 5’-RLM-RACE products at the different cleavage sites is shown in brackets. For amir296 transformed protoplasts, three colonies have the 5’-end at the expected position, opposite to nucleotides 10-11 of amir296, which is not the case for control and non-treated protoplasts.

### Predicted ***Si***sRNAs effectors are present in axenic cultures of *Serendipita indica*

After validation of the protoplast transformation strategy with artificial microRNAs, we next wanted to further characterize in the same system putative ck-*Si*sRNAs that we detected in our previous study in the *Si-Bd* symbiosis (Šečić et al., 2021). From the data set of 412 putative fungal ck-RNAs, we selected a set of 14 *Si*sRNA of 21 nt on the basis that they showed a higher expression level in colonized *Bd* roots, covered all four 5’-terminal nucleotides, and had interesting Arabidopsis predicted targets, involved in cell wall organisation, regulation of hormone signalling, immunity, and gene regulation. Using multiplexed stem-loop PCR followed by gel electrophoresis, we detected all 14 *Si*sRNAs in 4-week-old axenic *Si* culture (Figure 5). Next, we cloned a random subset of 6 out of 14 *Si*sRNAs (*Si*sRNA21, *Si*sRNA2*3*, *Si*sRNA24, *Si*sRNA28, *Si*sRNA154, and *Si*sRNA296) and subjected them to Sanger sequencing, which further confirmed the expected *Si*sRNA sequences.

**Figure 5:**
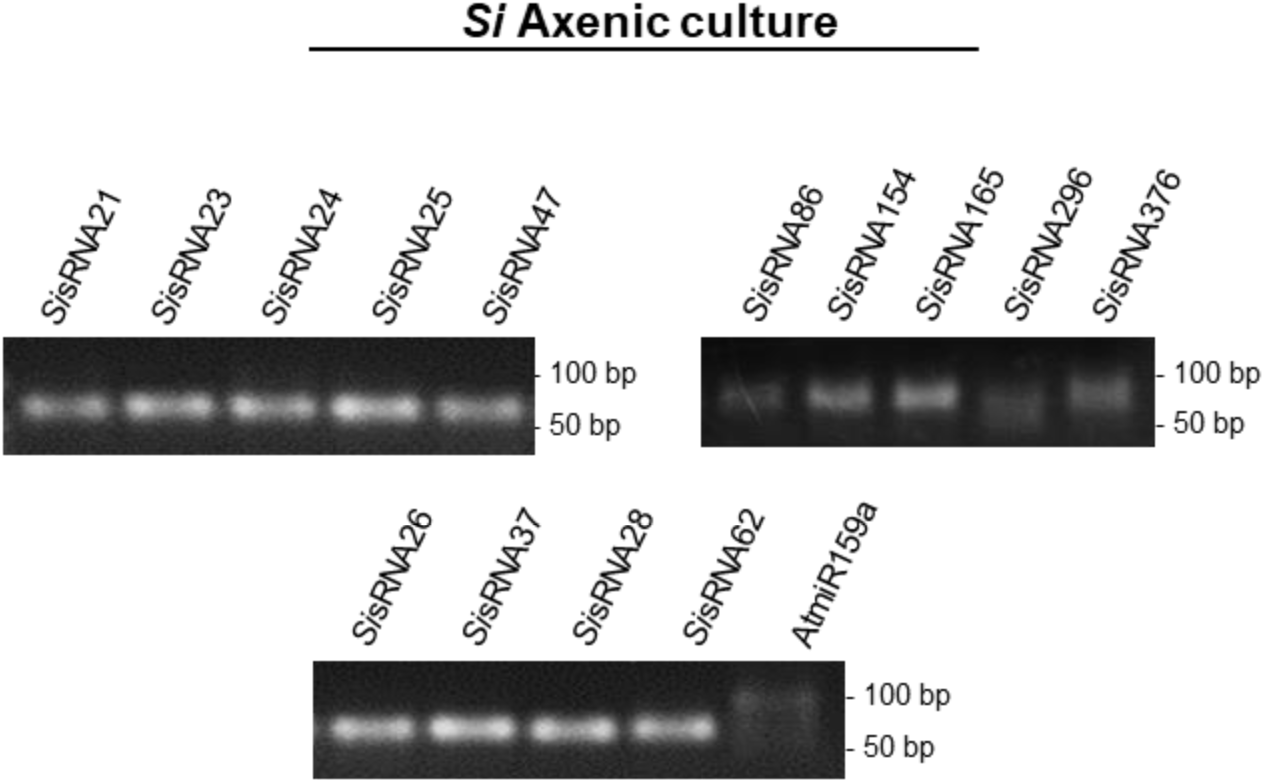
Gel electrophoresis of products from stem-loop PCR analysis of a set of potential SisRNA effectors from Serendipita indica. The *Si*RNAs were selected from previous RNAseq data sets generated from the mutualistic interaction of *Serendipita indica* and *Brachypodium distachyon* (Šečić et al., 2021). Multiplexed stem-loop PCR confirmed the presence of *Si*sRNAs in a 4-week-old *Si* axenic culture. A PCR product of the expected size of 62 bp (sum of the 21 bp of the sRNA sequence, loop region, plus the forward primer and the reverse primer length) is visualized in a 2% agarose gel. The plant *At*miR159a of 21 nt long served as a negative control. A weaker band of *Si*sRNA 296 was detected due to a lower abundance of the sRNA in the sample (see supplemental Table 8). An unspecific amplification (confirmed by Sanger sequencing) was detected for the *At*miR159a due to the multiplexing of hairpin primers or a possible partial complementarity of the hairpin primer to a non-target sequence.

### Predicted *Si*sRNAs effectors are present in colonized *Arabidopsis thaliana* roots

Next, we analysed the presence of a random subset of *Si*sRNAs (*Si*sRNA21, *Si*sRNA23, *Si*sRNA24, *Si*sRNA25, *Si*sRNA26, *Si*sRNA28, *Si*sRNA37, *Si*sRNA47, and *Si*sRNA62) (Supplemental Table 8) in *Si*-colonized *At* roots. Roots of 14-day-old seedlings were inoculated with chlamydospores and harvested at 3-, 7-, and 14 days post-inoculation (dpi). Fungal colonization of roots was confirmed after 7 days both by confocal laser scanning microscopy (CLSM) using the chitin-specific dye Wheat Germ Agglutinin with Alexa Fluor 488 (WGA-AF-488) (Figure 6 A) and by PCR using Internal Transcribed Spacer (Si*ITS)* primers and *Si*-specific *Ubiquitin* primers (Supplemental Figure 4). Multiplexed stem-loop PCR of total RNA extracted from *Si*-colonized *At* roots at different time points detected eight out of nine *Si*sRNAs (Figure 6 B). Bands for *Si*sRNA21 and *Si*sRNA24 did not consistently show up at all three-time points, likely because of the limitations associated with multiplexing the hairpin cDNA primers (Colombo et al., 2013; Kramer, 2011; Varkonyi-Gasic and Hellens, 2011). Consistent with this notion, the expression of *Si*sRNA21 and *Si*sRNA24 was detected at 7 dpi using stem-loop PCR with less multiplexing (Supplemental Figure 5). A random subset of the detected *Si*sRNAs (*Si*sRNA21, *Si*sRNA24, *Si*sRNA28, *Si*sRNA154) were cloned and submitted for Sanger sequencing, confirming the expected *Si*sRNA sequences in *Si*-colonized *At* roots.

**Figure 6:**
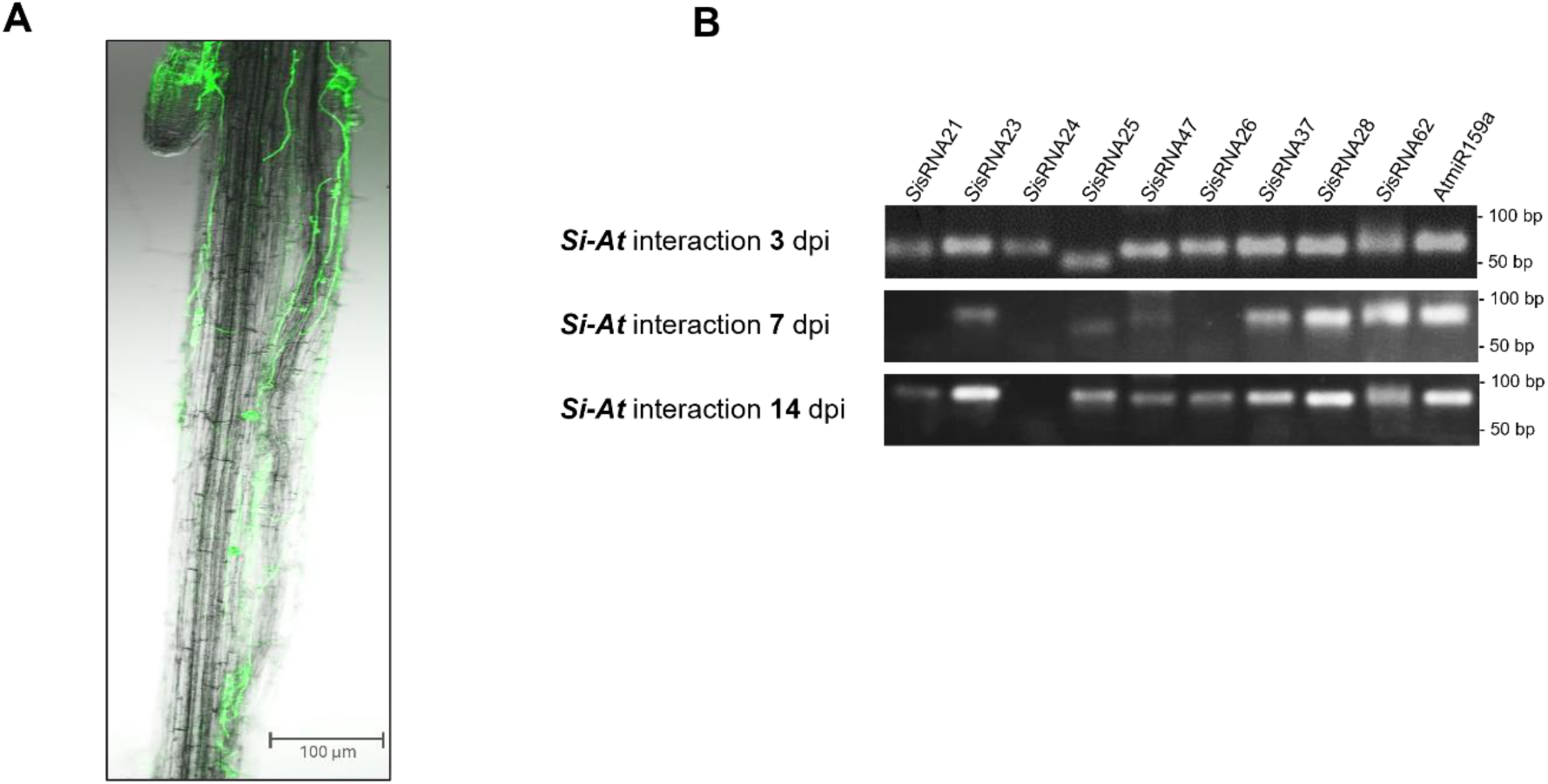
Detection of predicted *Si*sRNAs effectors in Arabidopsis roots colonized by *Serendipita indica*. (A) Root colonization pattern at 7 dpi. Fluorescence microscopy shows green chitin-specific WGA-AF488 staining of hyphal walls (λexc494 nm, λem515) (B) Detection of *Si*sRNAs in roots at 3-, 7- and 14 dpi using multiplexed stem-loop PCR. Agarose gel analysis of *Si*sRNA detected bands of the expected size of 62 bp. Plant *At*miR159a was used as a positive control for a plant-expressed sRNA.

### Predicted ***Si***sRNAs effectors mediate PTGS in transformed Arabidopsis protoplasts

Next, we transiently transformed protoplasts with *Si*sRNAs to demonstrate their gene-silencing activity. *At* protoplasts were transformed with a *35S::AtMIR390a* expression construct containing *Si*sRNA21, *Si*sRNA24, *Si*sRNA28 and *Si*sRNA154, respectively, (confirmed in *Si* axenic culture). Duplexed stem-loop PCR confirmed the expression and amplification of the four *Si*sRNAs. (Figure 7). The amplification products were cloned into the pGEM-T® cloning vector and sequencing confirmed their identity.

**Figure 7.**
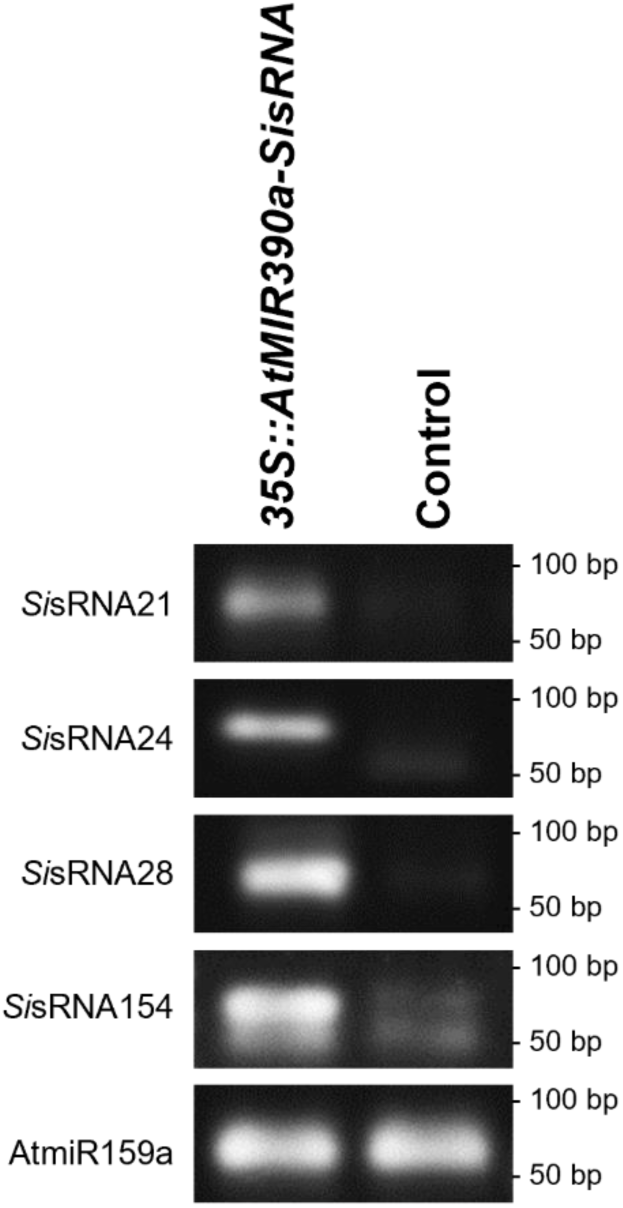
Duplexed stem-loop PCR confirming expression of *Si*sRNAs in Arabidopsis protoplasts. Agarose gel analysis of stem-loop PCR confirmed the detection of all four expressed *Si*sRNA in transformed protoplasts 24 hptr, but not in control protoplasts. Plant *At*miR159a was used as a positive control.

Next, we investigated the ability of the expressed *Si*sRNAs to induce PTGS of predicted target genes in protoplasts. *Si*sRNA24 was predicted to target five *At* genes: *AT1G57590 (PECTIN ACETYESTERASE 2)* involved in cell wall organization, *AT1G63180 (UDP-D-GALACTOSE 4-EPIMERASE 3)* involved in pollen development, *AT1G65090 (SEED LIPID DROPLET PROTEIN1*) involved in lipid droplet-plasma membrane tethering, *AT4G15765 (FAD/NAD(P)-binding oxidoreductase family protein)* involved in jasmonic acid-mediated signalling, and *AT5G16680* (PAIPP2, PHD2) involved in regulation of gene expression (see Supplemental File 2 and Supplemental Table 9). Expression of *Si*sRNA24, which has U at the 5’-terminus, and thus a preference for AGO1 (Mi *et al*., 2008; Montgomery *et al*., 2008; Takeda *et al*., 2008), significantly reduced the mRNA abundance of *AT1G63180*, *AT1G65090*, *AT4G15765* and *AT5G16680* in protoplasts 24 h after transformation by 63%, 56%, 45%, and 73%, respectively, (Figure 8 B). Similarly, *Si*sRNA154, also with 5’ U, is predicted to target *AT2G47600 (MAGNESIUM/PROTON EXCHANGER)* involved in magnesium, iron and zinc ion transport and *AT4G32160 (Phox (PX) domain-containing protein)* involved in signal transduction and previously investigated for PTGS in protoplasts transformed with amir154. Expression of *Si*sRNA154 significantly downregulated the accumulation of *AT2G47600* mRNA by 37% but not of *AT4G32160* (Figure 8 D). *Si*sRNA28 has 5’ Adenine (A), therefore a loading preference for AGO2 and AGO4 (Mi et al., 2008; Montgomery et al., 2008; Takeda et al., 2008). Expression of *Si*sRNA28 in protoplasts did not result in downregulation of any of the three predicted target genes: *AT1G05180 (AUXIN RESISTANT 1*) involved in auxin-activated signalling pathway and response to cytokinin, *AT5G55930* (*ARABIDOPSIS THALIANA OLIGOPEPTIDE TRANSPORTER 1*) an oligopeptide transporter, and *AT3G06670* (*PLATINUM SENSITIVE 2 LIKE*) with a regulatory function in ncRNA processing (Figure 8 C). Finally, expression of *Si*sRNA21 with 5’ Cytosine (C) and thus a preference for AGO5 (Mi et al., 2008; Montgomery et al., 2008; Takeda et al., 2008) resulted in downregulation of predicted target transcripts *AT5G25350* (a negative regulator of the ethylene-activated signalling pathway and previously investigated for PTGS in protoplasts transformed with ami*r*21), and *AT5G37600 (ARABIDOPSIS GLUTAMINE SYNTHASE 1)* involved in nitrate assimilation, by 51%, and 16%, respectively (Figure 8 A).

**Figure 8.**
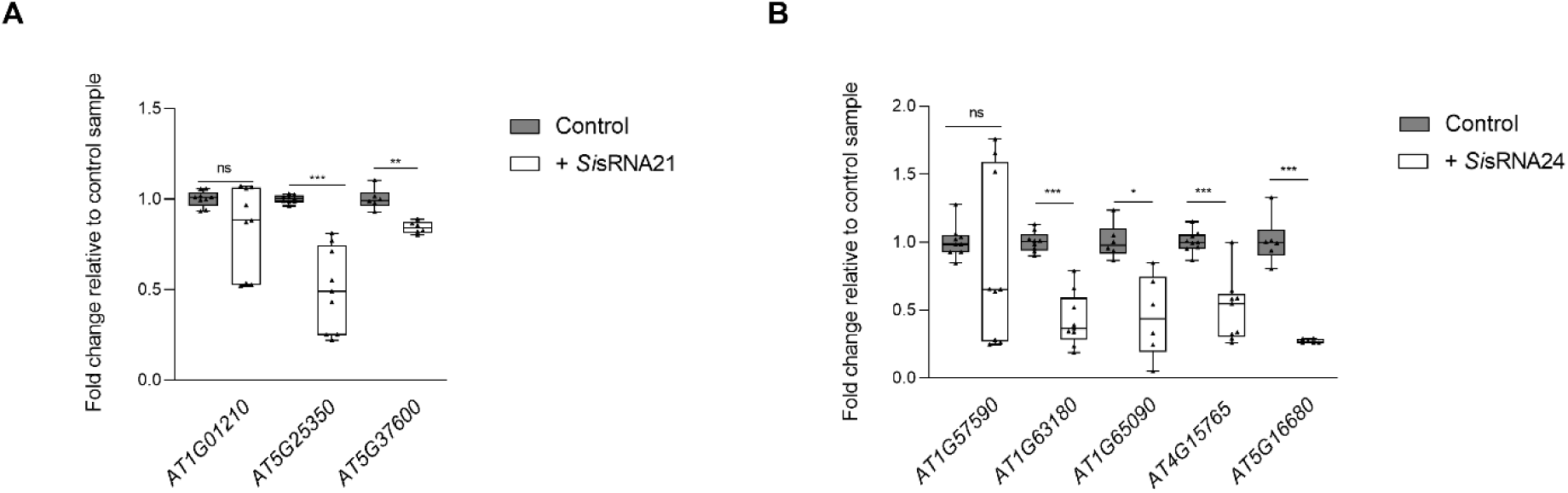
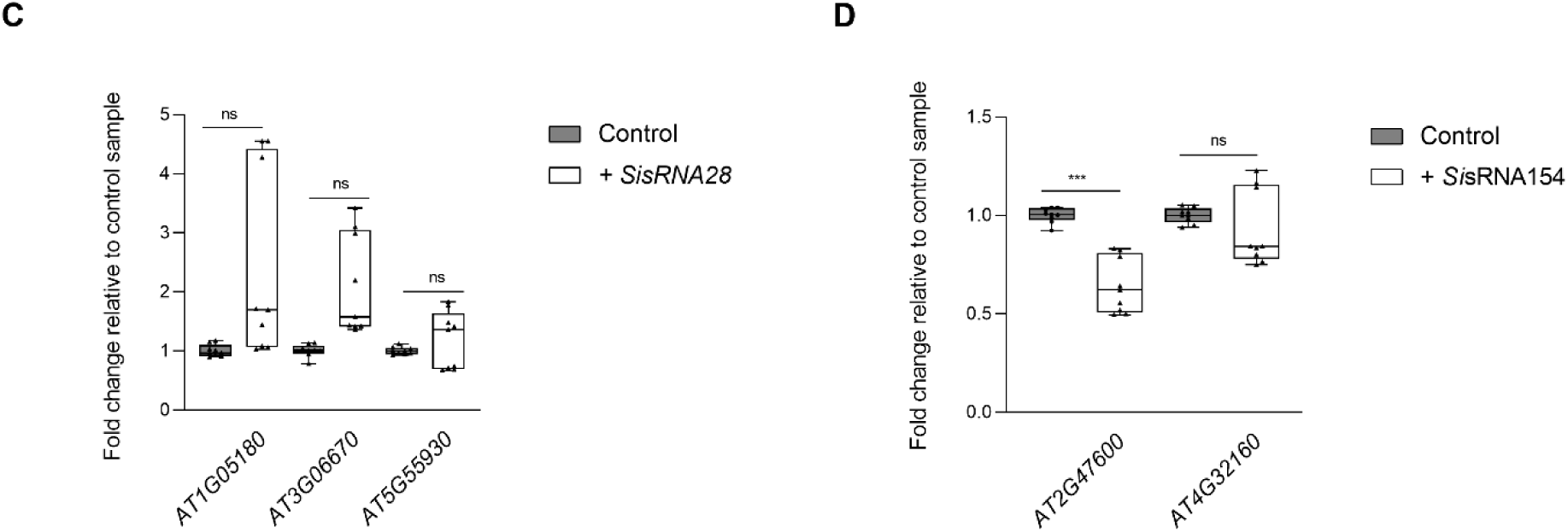
qPCR analysis of target gene silencing in Arabidopsis protoplasts 24 hptr with *Si*sRNAs vs. control (protoplast without SisRNA construct). The mRNA transcript levels in transformed protoplasts (+) expressing (A) *Si*sRNA21, (B) *Si*sRNA24, (C) *Si*sRNA28, and (D) *Si*sRNA154 normalized against housekeeping gene *Ubiquitin* (*AT3G62250*) are displayed as fold change relative to control protoplasts. Data are the average of two to three biological replicates ± standard deviation. Asterisks indicate difference at p ≤ 0.05(*), p ≤ 0.01 (**) and p ≤ 0.001 (***) according to Student’s t-test. ns= not significant.

### 5’-RLM-RACE reveals non-canonical cleavage of Arabidopsis target genes in protoplasts expressing SisRNAs with preference for AGO1

We selected *Si*sRNA24 to study the cleavage pattern of its significantly down-regulated target transcripts *AT1G65090* and *AT4G15765* using 5’-RLM-RACE. For the predicted target transcript *AT1G65090,* no band corresponding to an expected canonical cleavage product was detected (Supplemental Figure 6A). For target gene *AT4G15765*, we detected a single band of 300 bp in protoplasts expressing *Si*sRNA24, which was shorter than the expected size (443 bp), while double bands were observed for the control protoplasts. PCR amplicon from both transformed and control protoplasts were cloned into the pGEM-T^®^ cloning vector and sequenced. Sequence analysis confirmed the alignment of the 300 bp RLM-RACE product with *AT4G15765*. However, it did not confirm the 5’-end position, between nt 10 and 11, predicted for *Sis*RNA24-guided canonical cleavage (Supplemental Figure 6B). The absence of cleavage and/or a canonical cleavage product does not exclude the potential of *Si*sRNA24 in mediating predicted *At* gene silencing as many previous studies have highlighted the challenge of identifying sRNAs cleavage sites in plants (Brousse et al., 2014; Werner et al., 2021).

### Predicted ***Si***sRNAs effectors are loaded onto Arabidopsis AGO1

To further substantiate the hypothesis that ckRNAi occurs in the interaction between *Si* and *At* and that predicted *Si*sRNAs effectors are likely to be loaded into the Arabidopsis RISC complex, we conducted *At*AGO1/sRNA co-immunoprecipitation (Co-IP) using samples of *Si*-colonised roots, followed by stem-loop PCR analysis (Carbonell, 2017b; Dunker et al., 2021). To exclude the possibility that the *At*-specific anti-AGO1 antibody would cross-react with fungal AGOs, we first performed an *At*AGO1/sRNA Co-IP followed by Western-Blot (WB) analysis using a 4-week-old *Si* axenic culture. No binding of the *At*AGO1-specific antibody was detected in the crude extract neither in the supernatant nor in the IP fraction in comparison to the total loaded proteins from *Si* axenic culture confirming the specificity of the plant anti-AGO1 antibody (Supplementary Figure 7A). In line with this, no *Si*sRNAs could be detected by stem-loop PCR using *Si*sRNA-specific stem-loop primers (Supplementary Figure 7B). Similarly, western Blot analysis of *At*AGO1-IP of samples from *Si*-colonized *At* roots and *At* mock roots was performed. AGO1 accumulation was observed in crude extracts and IP fractions with a stronger signal in the IP fraction than in the supernatant from *Si-At* sample (Figures 9A and 9B).

**Figure 9.**
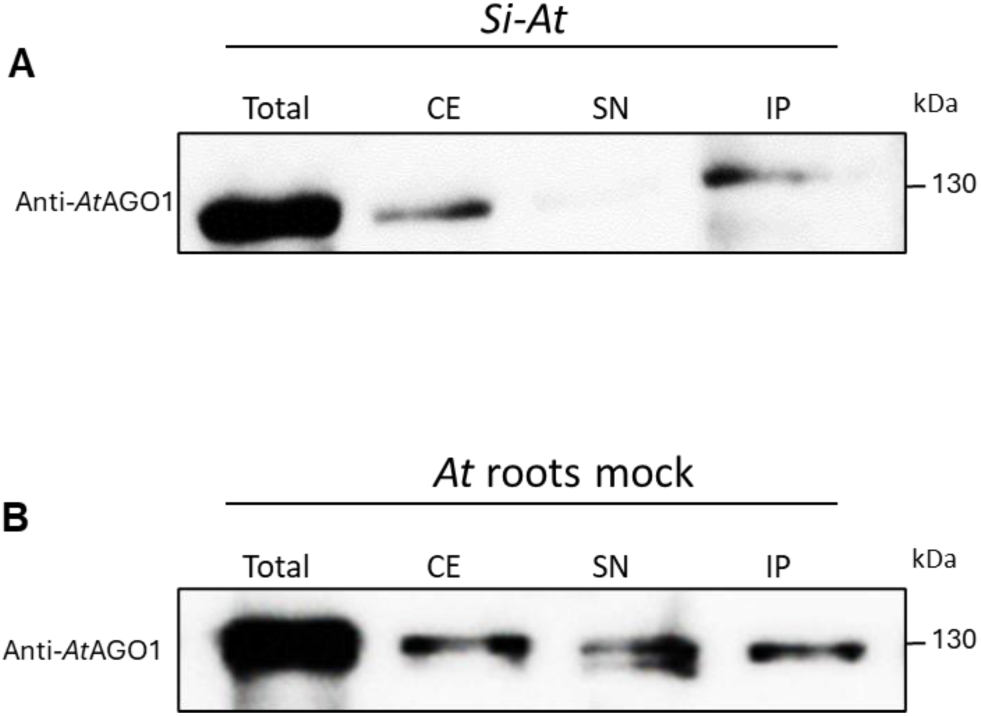
Quality control of AGO1 co-immunoprecipitation by western blot. Four sample fractions of the AGO1 co-IP experiment were analyzed: total protein before centrifugation and debris removal, crude extract (CE) after centrifugation and debris removal, supernatant (SN) after incubation with anti-*At*AGO1 and agarose beads, and IP fraction (resuspended IP pellet). These four fractions were analyzed in (A) *Si*-colonized root samples and (B) in *At* mock-treated roots. Both figures show the detection of *At*AGO1 using anti-*At*AGO1-specific antibody at the expected size of ⁓ 130 kDa. Note that AGO1 signals were stronger in IP fractions than in SN. The broad-range pre-stained protein marker was used as a protein-size marker.

Next, from *Si*-colonized roots (*Si-At*) Co-IP we recovered the *At*AGO1-bound sRNA sample. Stem-loop PCR was performed and detected the *At*AGO1-bound AtmiR159a and a band of the expected size for *Si*sRNA21, *Si*sRNA23, *Si*sRNA24, *Si*sRNA28 (Figure 10), whereas *At*miR393a*, which is known to preferably bind *At*AGO2 (Mi et al., 2008), was undetectable. Stem-loop PCR products from the *Si-At* Co-IP fraction were cloned and sequenced. The sequencing results confirmed the presence of plant *At*miR159a as well as *Si*sRNA2*4* and *Si*sRNA28 with the exact same sequences detected from *Si* axenic culture and *Si*-colonized *At* roots confirming the success of the pull-down. In addition, two *Si*sRNAs with almost identical sequences to *Si*sRNA21 and *Si*sRNA23 were also co-immunoprecipitated (Supplementary Table 10). Such mismatches that we detected for *Si*sRNA21 and *Si*sRNA23 may be caused by the processing steps of the sRNAs until the loading into the AGO1, which may result in sequence variations compared to the total RNAseq performed in our previous work (Šečić *et al*, 2021). Nevertheless, these results show that detected *Si*sRNAs are bound to *At*AGO1, supporting the notion that *Si*sRNAs are transferred into *At* root cells during root colonisation and loaded into the plant RNAi machinery.

**Figure 10:**
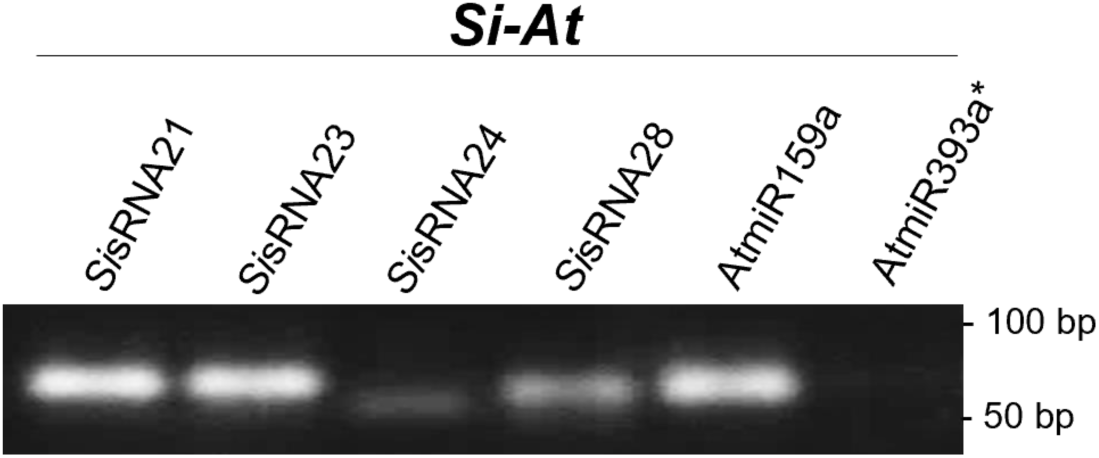
*Si*sRNAs co-immunoprecipitated with *At*AGO1 in Si-colonized Arabidopsis roots. Stem-loop PCR analysed in agarose gel confirms the presence of *Si*sRNA21, *Si*sRNA23, *Si*sRNA24, *Si*sRNA28, as well as AGO1 bound *At*miR159a but not AGO2 bound *At*miR393a* in *At*AGO1-co-IP samples from *Si*-colonized *At* roots.

## Discussion

In this work, we developed an experimental pipeline for the analysis of putative fungal sRNA effectors and investigated their role in host target gene silencing. We have previously reported the identification and analysis of fungal sRNAs induced in the interaction of the root endophyte *Serendipita indica* (*Si)* and *Brachypodium distachyon* (Šečić et al., 2021) termed ck-*Si*sRNAs. To bridge the gap between the large amount of bioinformatic data available from this previous study and the functional validation of the ck-*Si*sRNAs, we established here a rapid, simple and reproducible roadmap for the validation of ck-*Si*sRNA using Arabidopsis transient protoplast system. We first validated the effectiveness of our pipeline, by expressing artificial microRNAs (amiRNAs) and ck-*Si*sRNA in Arabidopsis protoplasts, confirmed their accumulation and their potential in inducing post-transcriptional gene silencing of the model plant *Arabidopsis thaliana* predicted target genes. Selected *Si*sRNAs were confirmed to be accumulated in *Si*-colonized roots and by immunoprecipitation pull-down assay, we showed that *Si*sRNAs are loaded into the plant RNAi machinery, confirming the translocation of *Si*sRNA effectors into host cells. Using *At*AGO/sRNA co-IP we detected the two *Si*sRNAs, *Si*sRNA24 and *Si*sRNA28 bound to *At*AGO1 supporting the ckRNAi hypothesis. Additional *Si*sRNAs of 21 nt were identified in the *At*AGO1 Co-IP assay that had some mismatches when compared to the *Si*sRNA detected in *Si* axenic culture and *Si-At* interaction.

To test the functionality of the selected ck-*Si*sRNAs in the *Si-At* symbiosis, we generated a plant expression construct under the 35S promotor for sRNA expression in protoplasts based on the pUC18 backbone vector and *At*MIR390a distal stem-loop which can generate 21 nt amiRNAs or *Si*sRNAs. The *At*MIR390a precursor belongs to the MIR390 family, which is highly conserved and precisely produced and processed in several plant species (Axtell et al., 2006; Cuperus et al., 2011). Carbonell et al (2014) tested the functionality of the *At*MIR390a-based amiRNAs vector in repressing target transcript accumulation in Arabidopsis plants using the plant vector pMDC32BAtMIR390a-B/c, which enables the directed cloning of amiRNAs and their stable expression in dicotyledonous plants (Carbonell et al., 2014). Although MIR390 associates preferably with AGO7, the association of *At*MIR390a-derived amiRNAs that have a 5’ U can be directed to AGO1 (Carbonell et al., 2014; Carbonell, 2017b; Cuperus et al., 2010; Montgomery et al., 2008). Conventionally, an sRNA-AGO1-mediated cleavage results in a canonical cut between nucleotides 10 and 11 of the sRNA at the corresponding mRNA site. This cleavage is a confirmation of sRNA loading into AGO1, formation of the RISC complex and degradation of the mRNA target (Addo-Quaye *et al*., 2009; Huntzinger & Izaurralde, 2011). In our approach, we combined the advantages of plant expression vectors and protoplast transient expression systems, both considered as valuable tools, to study gene silencing and the molecular mechanisms of RNAi in plants (Tyurin et al., 2020). To confirm the effectiveness of the transient protoplasts system, we first used artificial microRNAs (amiRNAs) with almost a perfect match to *At’s* predicted target transcript. Four amiRNAs, amir21, amir24, amir154 and amir296 (Supplemental Table 2, were expressed in *At* protoplasts. All amiRNAs induced a significant reduction in mRNA abundance of their corresponding host-target transcripts, confirming their effective silencing activity in *At* protoplasts. AmiRNAs used in our pipeline are algorithmically designed to exhibit a 5’ U, thereby enabling their loading into the AGO1 protein (Carbonell, 2017c). Consequently, they can enter the RISC complex and induce a downregulation of the target gene. We then investigated whether amir296 that showed a reduction of 65% of the predicted target *AT2G45240* (MAP1A, METHIONINE AMINOPEPTIDASE 1A), can mediate a canonical cleavage of the same target *AT2G45240,* and indeed detected using 5’-RLM-RACE a cleavage site of the mRNA transcript between position 10 and 11 of amir296 indicating that it can induce a potential *At*AGO1-mediated degradation of *AT2G45240* mRNA.

Next, we expressed a set of the selected putative ck-*Si*sRNAs in Arabidopsis protoplasts. The expression and accumulation of the putative *Si*sRNA24, exhibiting a 5’-U, was validated by stem-loop PCR. Four of five predicted targets of *Si*sRNA24 showed a significantly reduced abundance in the transformed protoplasts. Another putative *SisRNA*, *SisRNA154*, which also has a 5’-U, showed significant downregulation only for the target *AT2G47600* from the two *At* predicted target genes.

An important factor to consider when studying PTGS via sRNA effectors is the degree of mismatches, expressed as “Expectation value” reporting on the mismatch score between the predicted target mRNA and the sRNA. Previous studies have demonstrated that the number and position of mismatches between the sRNA and the corresponding target mRNA can significantly impact their binding affinity, eventually weakening the interaction and resulting in either non-significant or no downregulation (Carbonell et al., 2015; Ossowski et al., 2008; Rhoades et al., 2002). Studies have shown that mismatches within the sRNA seed region spanning nt 2 to 12/13 are possible, but they may reduce the sRNA activity, whereas mismatches at position 1 or between nt 14 and 21 are more acceptable (Carbonell, 2017a; Mallory *et al*., 2004). Moreover, mismatches can also lead to a different mode of action for the sRNA, and instead of an mRNA degradation pathway, mismatches can trigger a deadenylation pathway or a translational repression pathway, thus causing a decrease in protein synthesis instead of an mRNA target degradation (Baulcombe, 2004; Carthew & Sontheimer, 2009). We also hypothesize that a high abundance level of a target mRNA may lead to less detectable downregulation, as its accessibility for the sRNA may have an impact on the silencing process and the detection of a canonical cleavage. It is also conceivable that the presence of other endogenous miRNAs, which may bind near or at the same mRNA binding site might disrupt sRNA/mRNA binding, modulate the regulatory outcome, and affect the gene expression pattern.

Further analysis of the molecular cleavage pattern of *Si*sRNA24 by 5’-RLM-RACE identified noncanonical cut sites in the target gene *AT4G15765.* This finding may be explained by the fact that cleavage products might be present at a low abundance, falling below the detection threshold of the RLM-RACE method and making the detection of a precise canonical cleavage a challenge. Alternatively, it might be also possible that the cleavage site within the target mRNA could be located at a different position. Moreover, 5’-RLM-RACE is a challenging method to detect cleavage sites. High throughput sequencing of samples from the *Si-At* interaction including transcriptional analysis and degradome sequencing are needed to better understand *Si*sRNA-mediated target cleavage. Hence, consideration for sRNA non-canonical cleavage site may be further investigated as different studies speculate that PTGS can still be induced through sRNA or phasiRNA and generate a non-canonical cleavage site (Brousse et al., 2014; Ivanova et al., 2022).

In our study, the Arabidopsis target *AT5G25350* is predicted to be targeted by both amir21 and *Si*sRNA21. When expressing amir21 (with 5’ U and an expectation value of 0), the mRNA abundance of *AT5G25350* showed a downregulation of 68%. However, when expressing *Si*sRNA21 (with a 5’C and an expectation value of 5) we observed a downregulation of 51% of the same target gene. Another example is the predicted target gene *AT4G32160* targeted by amir154 and *Si*sRNA154 both with a 5’ U. The mRNA abundance of *AT4G32160* in protoplasts expressing amir154 (with an expectation value of 0) resulted in a reduction of 50% but protoplasts expressing *Si*sRNA154 (with an expectation value of 4.5) did not show a reduction of *AT4G32160* transcript (Supplemental Figure 8). These results raise the possibility that amiRNAs are more prone to mediate target gene silencing but further studies with additional amiRNAs and putative sRNAs are required to confirm these observations.

We extended our investigation by performing *At*AGO1/sRNA Co-IP on samples from *Si*-colonized Arabidopsis roots and western-blot analysis confirmed the enrichment of the *At*AGO1 in the IP fraction. We subsequently detected using stem-loop PCR two ck-*Si*sRNAs: *Si*sRNA24 and *Si*sRNA28, bound to *At*AGO1 providing strong evidence of the translocation of *Si*sRNAs and their incorporation into Arabidopsis RNAi machinery. Importantly, the identification in the *At*AGO1-IP of two *Si*sRNAs exhibiting nucleotide variations at their 3’terminal end to the previously identified ck-*Si*sRNA (*Si*sRNA21 and *Si*sRNA23) suggests alternative *Si*sRNA variants. The presence of such variants highlights the complexity of the sRNA-mediated regulatory networks during *Si*-host root colonization. Conducting a high throughput sequencing of the entire pool of *Si*sRNA pulled down from *At*AGO1 Co-IP could identify a large “Repertoire” of *Si*sRNAs effectors with a functional role in *Si* symbiosis.

In conclusion, we have developed a transient protoplast expression system to investigate the potential role of putative *Si*sRNA effectors in regulating target gene expression in *Arabidopsis thaliana*. This validation tool will not only help to understand the underlying molecular mechanism of *Si*sRNA-mediated PTGS but will also allow us to examine ckRNAi in mutualistic plant-fungal association.

By studying *Si*sRNA-mediated PTGS, we have gained first insights into the possibility that the endophytic fungus *Si* exploits sRNA effectors for the establishment of its mutualistic symbiosis with plants. More targeted studies will confirm whether *Si* utilizes its sRNAs to modulate the plant immune system and identify more genes that can be silenced during the *Si-At* interaction. With the great potential of *Si* to colonize a wide host range of plants, we expect that findings from the study of the *Si-At* interaction can be translated to crop plants. Moreover, amiRNAs have been shown to be effective in fine-tuning gene expression in plants with a high silencing level (Carbonell et al., 2014; Cisneros & Carbonell, 2020). Therefore, they can be viewed as a potential tool for RNAi-mediated crop protection strategies. Computational tools, along with *in vivo* and *in vitro* experimental setups, can expand our knowledge of ckRNAi and reveal the unknown about mutualistic and pathogenic interaction for cost-effective basic research and agronomic application.

## Abbreviations

AGO: argonaute
*At*: *Arabidopsis thaliana*
DCL: dicer-like
dpi: days post inoculation
hptr: hours post-transformation
IP: immunoprecipitation
miRNA: micro RNA
nt: nucleotide
PTGS: post-transcriptional gene silencing
RISC: RNA-induced silencing complex
RKN: root-knot nematodes
RNAi: RNA interference
*Si*: *Serendipita indica*
siRNA: small interfering RNA
sRNA: small RNA

## Acknowledgments

The authors gratefully thank R. Eichmann for critically reading and correcting the manuscript. We thank M. Ladera-Carmona and B.T. Werner for their comments and suggestions. We thank the German Research Foundation (DFG) for supporting our research on RNA in plant-pathogen interactions by granting the DFG research unit RU5116 on exRNA to KHK. SN is supported by the Dr. Ernst-Leopold Klipstein-Stiftung, Paderborn, Giessen.

## Conflict of interest

The authors declare that they have no competing interests.

## Author contributions

S.N. and J.S. conceived the study; S.N. and K-H.K. wrote the paper; J.S. and E. Š. generated prediction of Arabidopsis targets from sRNAseq data. S.N., K.B., and S.S. conducted the functional experiments. All authors read and approved the final manuscript.

## Notes

### Competing Interest Statement

The authors have declared no competing interest.

### Summary of Updates

SisRNA selection and Arabidopsis target prediction analysis Developed AGO1 immunoprecipitation assay including Western-blot analysis Supplemental files updated

## References

Addo-Quaye, C., Snyder, J. A., Park, Y. B., Li, Y.-F., Sunkar, R., & Axtell, M. J. (2009). Sliced microRNA targets and precise loop-first processing of MIR319 hairpins revealed by analysis of the Physcomitrella patens degradome. RNA, 15(12), 2112–2121. 10.1261/rna.1774909

Adhikari, S., Turner, M., & Subramanian, S. (2013). Hairpin Priming Is Better Suited than In Vitro Polyadenylation to Generate cDNA for Plant miRNA qPCR. Molecular Plant, 6(1), 229–231. 10.1093/mp/sss106

Akum, F. N., Steinbrenner, J., Biedenkopf, D., Imani, J., & Kogel, K.-H. (2015). The Piriformospora indica effector PIIN_08944 promotes the mutualistic Sebacinalean symbiosis. Frontiers in Plant Science, 6, 906. 10.3389/fpls.2015.00906

Axtell, M. J., Jan, C., Rajagopalan, R., & Bartel, D. P. (2006). A Two-Hit Trigger for siRNA Biogenesis in Plants. Cell, 127(3), 565–577. 10.1016/j.cell.2006.09.032

Bargmann, B. O. R., & Birnbaum, K. D. (2009). Positive Fluorescent Selection Permits Precise, Rapid, and In-Depth Overexpression Analysis in Plant Protoplasts. Plant Physiology, 149(3), 1231– 1239. 10.1104/pp.108.133975

Baulcombe, D. (2004). RNA silencing in plants. Nature, 431(7006), Article 7006. 10.1038/nature02874

Brousse, C., Liu, Q., Beauclair, L., Deremetz, A., Axtell, M. J., & Bouché, N. (2014). A non-canonical plant microRNA target site. Nucleic Acids Research, 42(8), 5270–5279. 10.1093/nar/gku157

Cai, Q., Qiao, L., Wang, M., He, B., Lin, F.-M., Palmquist, J., Huang, S.-D., & Jin, H. (2018). Plants send small RNAs in extracellular vesicles to fungal pathogen to silence virulence genes. Science (New York, N.Y.), 360(6393), 1126–1129. 10.1126/science.aar4142

Carbonell, A. (2017a). Artificial small RNA-based strategies for effective and specific gene silencing in plants. Plant Gene Silencing: Mechanisms and Applications, 110–127. 10.1079/9781780647678.0110

Carbonell, A. (2017b). Immunoprecipitation and High-Throughput Sequencing of ARGONAUTE-Bound Target RNAs from Plants. In A. Carbonell (Ed.), Plant Argonaute Proteins: Methods and Protocols (pp. 93–112). Springer. 10.1007/978-1-4939-7165-7_6

Carbonell, A. (2017c). Plant ARGONAUTEs: Features, Functions, and Unknowns. In A. Carbonell (Ed.), Plant Argonaute Proteins: Methods and Protocols (pp. 1–21). Springer. 10.1007/978-1-4939-7165-7_1

Carbonell, A., Fahlgren, N., Mitchell, S., Cox Jr, K., Mockler, T., & Carrington, J. (2015). Highly Specific Gene Silencing in a Monocot Species by Artificial MicroRNAs Derived From Chimeric MIRNA Precursors. Plant J. (in Press). 10.1111/tpj.12835

Carbonell, A., Takeda, A., Fahlgren, N., Johnson, S. C., Cuperus, J. T., & Carrington, J. C. (2014). New generation of artificial MicroRNA and synthetic trans-acting small interfering RNA vectors for efficient gene silencing in Arabidopsis. Plant Physiology, 165(1), 15–29. 10.1104/pp.113.234989

Carthew, R. W., & Sontheimer, E. J. (2009). Origins and Mechanisms of miRNAs and siRNAs. Cell, 136(4), 642–655. 10.1016/j.cell.2009.01.035

Cisneros, A. E., & Carbonell, A. (2020). Artificial Small RNA-Based Silencing Tools for Antiviral Resistance in Plants. Plants, 9(6), Article 6. 10.3390/plants9060669

Colombo, S., Nielsen, H. M., & Foged, C. (2013). Evaluation of carrier-mediated siRNA delivery: Lessons for the design of a stem-loop qPCR-based approach for quantification of intracellular full-length siRNA. Journal of Controlled Release, 166(3), 220–226. 10.1016/j.jconrel.2013.01.006

Cuperus, J. T., Carbonell, A., Fahlgren, N., Garcia-Ruiz, H., Burke, R. T., Takeda, A., Sullivan, C. M., Gilbert, S. D., Montgomery, T. A., & Carrington, J. C. (2010). Unique Functionality of 22 nt miRNAs in Triggering RDR6-Dependent siRNA Biogenesis from Target Transcripts in Arabidopsis. Nature Structural & Molecular Biology, 17(8), 997–1003. 10.1038/nsmb.1866

Cuperus, J. T., Fahlgren, N., & Carrington, J. C. (2011). Evolution and Functional Diversification of MIRNA Genes. The Plant Cell, 23(2), 431–442. 10.1105/tpc.110.082784

Dai, X., Zhuang, Z., & Zhao, P. X. (2018). psRNATarget: A plant small RNA target analysis server (2017 release). Nucleic Acids Research, 46(Web Server issue), W49–W54. 10.1093/nar/gky316

Dunker, F., Lederer, B., & Weiberg, A. (2021). Plant ARGONAUTE Protein Immunopurification for Pathogen Cross Kingdom Small RNA Analysis. Bio-Protocol, 11(3), e3911. 10.21769/BioProtoc.3911

Fahlgren, N., Hill, S. T., Carrington, J. C., & Carbonell, A. (2016). P-SAMS: A web site for plant artificial microRNA and synthetic trans-acting small interfering RNA design. Bioinformatics, 32(1), 157–158. 10.1093/bioinformatics/btv534

Glaeser, S. P., Imani, J., Alabid, I., Guo, H., Kumar, N., Kämpfer, P., Hardt, M., Blom, J., Goesmann, A., Rothballer, M., Hartmann, A., & Kogel, K.-H. (2016). Non-pathogenic Rhizobium radiobacter F4 deploys plant beneficial activity independent of its host Piriformospora indica. The ISME Journal, 10(4), 871–884. 10.1038/ismej.2015.163

Göhre, V., & Weiberg, A. (2023). RNA Dialogues in Fungal–Plant Relationships. In B. Scott & C. Mesarich (Eds.), Plant Relationships: Fungal-Plant Interactions (pp. 31–51). Springer International Publishing. 10.1007/978-3-031-16503-0_2

He, B., Cai, Q., Qiao, L., Huang, C.-Y., Wang, S., Miao, W., Ha, T., Wang, Y., & Jin, H. (2021). RNA-binding proteins contribute to small RNA loading in plant extracellular vesicles. Nature Plants, 7(3), 342–352. 10.1038/s41477-021-00863-8

He, B., Hamby, R., & Jin, H. (2021). Plant extracellular vesicles: Trojan horses of cross-kingdom warfare. FASEB BioAdvances, 3(9), 657–664. 10.1096/fba.2021-00040

Huang, G., Allen, R., Davis, E. L., Baum, T. J., & Hussey, R. S. (2006). Engineering broad root-knot resistance in transgenic plants by RNAi silencing of a conserved and essential root-knot nematode parasitism gene. Proceedings of the National Academy of Sciences, 103(39), 14302–14306. 10.1073/pnas.0604698103

Huntzinger, E., & Izaurralde, E. (2011). Gene silencing by microRNAs: Contributions of translational repression and mRNA decay. Nature Reviews Genetics, 12(2), Article 2. 10.1038/nrg2936

Ivanova, Z., Minkov, G., Gisel, A., Yahubyan, G., Minkov, I., Toneva, V., & Baev, V. (2022). The Multiverse of Plant Small RNAs: How Can We Explore It? International Journal of Molecular Sciences, 23(7), 3979. PubMed. 10.3390/ijms23073979

Iwakawa, H., & Tomari, Y. (2022). Life of RISC: Formation, action, and degradation of RNA-induced silencing complex. Molecular Cell, 82(1), 30–43. 10.1016/j.molcel.2021.11.026

Koch, A., Kumar, N., Weber, L., Keller, H., Imani, J., & Kogel, K.-H. (2013). Host-induced gene silencing of cytochrome P450 lanosterol C14α-demethylase–encoding genes confers strong resistance to Fusarium species. Proceedings of the National Academy of Sciences, 110(48), 19324– 19329. 10.1073/pnas.1306373110

Kramer, M. F. (2011). STEM-LOOP RT-qPCR for miRNAS. *Current Protocols in Molecular Biology / Edited by Frederick M. Ausubel … [et Al.]*, *CHAPTER*, Unit15.10. 10.1002/0471142727.mb1510s95

Lincoln, C., Britton, J. H., & Estelle, M. (1990). Growth and development of the axr1 mutants of Arabidopsis. The Plant Cell, 2(11), 1071–1080.

Liu, S., Jaouannet, M., Dempsey, D. A., Imani, J., Coustau, C., & Kogel, K.-H. (2020). RNA-based technologies for insect control in plant production. Biotechnology Advances, 39, 107463. 10.1016/j.biotechadv.2019.107463

Livak, K. J., & Schmittgen, T. D. (2001). Analysis of relative gene expression data using real-time quantitative PCR and the 2(-Delta Delta C(T)) Method. Methods (San Diego, Calif.), 25(4), 402–408. 10.1006/meth.2001.1262

Llave, C., Xie, Z., Kasschau, K. D., & Carrington, J. C. (2002). Cleavage of Scarecrow-like mRNA Targets Directed by a Class of Arabidopsis miRNA. Science, 297(5589), 2053–2056. 10.1126/science.1076311

Mahanty, B., Mishra, R., & Joshi, R. K. (2023). Cross-kingdom small RNA communication between plants and fungal phytopathogens-recent updates and prospects for future agriculture. RNA Biology, 20(1), 109–119. 10.1080/15476286.2023.2195731

Mallory, A. C., Reinhart, B. J., Jones-Rhoades, M. W., Tang, G., Zamore, P. D., Barton, M. K., & Bartel, D. P. (2004). MicroRNA control of PHABULOSA in leaf development: Importance of pairing to the microRNA 5′ region. The EMBO Journal, 23(16), 3356–3364. 10.1038/sj.emboj.7600340

Meyers, B. C., & Axtell, M. J. (2019). MicroRNAs in Plants: Key Findings from the Early Years. The Plant Cell, 31(6), 1206–1207. 10.1105/tpc.19.00310

Mi, S., Cai, T., Hu, Y., Chen, Y., Hodges, E., Ni, F., Wu, L., Li, S., Zhou, H., Long, C., Chen, S., Hannon, G. J., & Qi, Y. (2008). Sorting of Small RNAs into Arabidopsis Argonaute Complexes Is Directed by the 5′ Terminal Nucleotide. Cell, 133(1), 116–127. 10.1016/j.cell.2008.02.034

Montgomery, T. A., Howell, M. D., Cuperus, J. T., Li, D., Hansen, J. E., Alexander, A. L., Chapman, E. J., Fahlgren, N., Allen, E., & Carrington, J. C. (2008). Specificity of ARGONAUTE7-miR390 interaction and dual functionality in TAS3 trans-acting siRNA formation. Cell, 133(1), 128–141. 10.1016/j.cell.2008.02.033

Moreno, P., Fexova, S., George, N., Manning, J. R., Miao, Z., Mohammed, S., Muñoz-Pomer, A., Fullgrabe, A., Bi, Y., Bush, N., Iqbal, H., Kumbham, U., Solovyev, A., Zhao, L., Prakash, A., García-Seisdedos, D., Kundu, D. J., Wang, S., Walzer, M., … Papatheodorou, I. (2022). Expression Atlas update: Gene and protein expression in multiple species. Nucleic Acids Research, 50(D1), D129–D140. 10.1093/nar/gkab1030

Nasfi, S., & Kogel, K.-H. (2022). Packaged or unpackaged: Appearance and transport of extracellular noncoding RNAs in the plant apoplast. *ExRNA; Vol 4 (May 31, 2022): ExRNA*. https://exrna.amegroups.com/article/view/64855

Nowara, D., Gay, A., Lacomme, C., Shaw, J., Ridout, C., Douchkov, D., Hensel, G., Kumlehn, J., & Schweizer, P. (2010). HIGS: Host-Induced Gene Silencing in the Obligate Biotrophic Fungal Pathogen *Blumeria graminis*. The Plant Cell, 22(9), 3130–3141. 10.1105/tpc.110.077040

Osborne, R., Rehneke, L., Lehmann, S., Roberts, J., Altmann, M., Altmann, S., Zhang, Y., Köpff, E., Dominguez-Ferreras, A., Okechukwu, E., Sergaki, C., Rich-Griffin, C., Ntoukakis, V., Eichmann, R., Shan, W., Falter-Braun, P., & Schäfer, P. (2023). Symbiont-host interactome mapping reveals effector-targeted modulation of hormone networks and activation of growth promotion. Nature Communications, 14(1), Article 1. 10.1038/s41467-023-39885-5

Ossowski, S., Schwab, R., & Weigel, D. (2008). Gene silencing in plants using artificial microRNAs and other small RNAs. The Plant Journal, 53(4), 674–690. 10.1111/j.1365-313X.2007.03328.x

Qiang, X., Weiss, M., Kogel, K.-H., & Schäfer, P. (2012). Piriformospora indica—A mutualistic basidiomycete with an exceptionally large plant host range. Molecular Plant Pathology, 13(5), 508–518. 10.1111/j.1364-3703.2011.00764.x

Qiao, S. A., Gao, Z., & Roth, R. (2023). A perspective on cross-kingdom RNA interference in mutualistic symbioses. New Phytologist, 240(1), 68–79. 10.1111/nph.19122

Reinhart, B. J., Weinstein, E. G., Rhoades, M. W., Bartel, B., & Bartel, D. P. (2002). MicroRNAs in plants. Genes & Development, 16(13), 1616–1626. 10.1101/gad.1004402

Ren, B., Wang, X., Duan, J., & Ma, J. (2019). Rhizobial tRNA-derived small RNAs are signal molecules regulating plant nodulation. Science (New York, N.Y.), 365(6456), 919–922. 10.1126/science.aav8907

Rhoades, M. W., Reinhart, B. J., Lim, L. P., Burge, C. B., Bartel, B., & Bartel, D. P. (2002). Prediction of Plant MicroRNA Targets. Cell, 110(4), 513–520. 10.1016/S0092-8674(02)00863-2

Rutter, B. D., & Innes, R. W. (2018). Extracellular vesicles as key mediators of plant–microbe interactions. Current Opinion in Plant Biology, 44, 16–22. 10.1016/j.pbi.2018.01.008

Sang, H., & Kim, J.-I. (2020). Advanced strategies to control plant pathogenic fungi by host-induced gene silencing (HIGS) and spray-induced gene silencing (SIGS). Plant Biotechnology Reports, 14(1), 1–8. 10.1007/s11816-019-00588-3

Šečić, E., Zanini, S., Wibberg, D., Jelonek, L., Busche, T., Kalinowski, J., Nasfi, S., Thielmann, J., Imani, J., Steinbrenner, J., & Kogel, K.-H. (2021). A novel plant-fungal association reveals fundamental sRNA and gene expression reprogramming at the onset of symbiosis. BMC Biology, 19(1), 171. 10.1186/s12915-021-01104-2

Silvestri, A., Fiorilli, V., Miozzi, L., Accotto, G. P., Turina, M., & Lanfranco, L. (2019). In silico analysis of fungal small RNA accumulation reveals putative plant mRNA targets in the symbiosis between an arbuscular mycorrhizal fungus and its host plant. BMC Genomics, 20, 169. 10.1186/s12864-019-5561-0

Takeda, A., Iwasaki, S., Watanabe, T., Utsumi, M., & Watanabe, Y. (2008). The Mechanism Selecting the Guide Strand from Small RNA Duplexes is Different Among Argonaute Proteins. Plant and Cell Physiology, 49(4), 493–500. 10.1093/pcp/pcn043

Tyurin, A. A., Suhorukova, A. V., Kabardaeva, K. V., & Goldenkova-Pavlova, I. V. (2020). Transient Gene Expression is an Effective Experimental Tool for the Research into the Fine Mechanisms of Plant Gene Function: Advantages, Limitations, and Solutions. Plants, 9(9), 1187. 10.3390/plants9091187

Ueno, D., Yamasaki, S., & Kato, K. (2022). Methods for detecting RNA degradation intermediates in plants. Plant Science, 318, 111241. 10.1016/j.plantsci.2022.111241

Varkonyi-Gasic, E., & Hellens, R. P. (2011). Quantitative stem-loop RT-PCR for detection of microRNAs. Methods in Molecular Biology (Clifton, N.J.), 744, 145–157. 10.1007/978-1-61779-123-9_10

Verma, N., Narayan, O. P., Prasad, D., Jogawat, A., Panwar, S. L., Dua, M., & Johri, A. K. (2022). Functional characterization of a high-affinity iron transporter (PiFTR) from the endophytic fungus Piriformospora indica and its role in plant growth and development. Environmental Microbiology, 24(2), 689–706. 10.1111/1462-2920.15659

Verma, S., Varma, A., Rexer, K.-H., Hassel, A., Kost, G., Sarbhoy, A., Bisen, P., Bütehorn, B., & Franken, P. (1998). Piriformospora indica, gen. Et sp. Nov., a new root-colonizing fungus. Mycologia, 90(5), 896–903. 10.1080/00275514.1998.12026983

Wang, M., & Dean, R. A. (2020). Movement of small RNAs in and between plants and fungi. Molecular Plant Pathology, 21(4), 589–601. 10.1111/mpp.12911

Wang, M., Thomas, N., & Jin, H. (2017). Cross-kingdom RNA trafficking and environmental RNAi for powerful innovative pre- and post-harvest plant protection. Current Opinion in Plant Biology, 38, 133–141. 10.1016/j.pbi.2017.05.003

Weiberg, A., Wang, M., Lin, F.-M., Zhao, H., Zhang, Z., Kaloshian, I., Huang, H.-D., & Jin, H. (2013). Fungal small RNAs suppress plant immunity by hijacking host RNA interference pathways. Science (New York, N.Y.), 342(6154), 118–123. 10.1126/science.1239705

Weiß, M., Waller, F., Zuccaro, A., & Selosse, M.-A. (2016). Sebacinales – one thousand and one interactions with land plants. New Phytologist, 211(1), 20–40. 10.1111/nph.13977

Werner, B. T., Koch, A., Šečić, E., Engelhardt, J., Jelonek, L., Steinbrenner, J., & Kogel, K.-H. (2021). Fusarium graminearum DICER-like-dependent sRNAs are required for the suppression of host immune genes and full virulence. PLOS ONE, 16(8), e0252365. 10.1371/journal.pone.0252365

Wong-Bajracharya, J., Singan, V. R., Monti, R., Plett, K. L., Ng, V., Grigoriev, I. V., Martin, F. M., Anderson, I. C., & Plett, J. M. (2022). The ectomycorrhizal fungus Pisolithus microcarpus encodes a microRNA involved in cross-kingdom gene silencing during symbiosis. Proceedings of the National Academy of Sciences of the United States of America, 119(3), e2103527119. 10.1073/pnas.2103527119

Wu, F.-H., Shen, S.-C., Lee, L.-Y., Lee, S.-H., Chan, M.-T., & Lin, C.-S. (2009). Tape-Arabidopsis Sandwich—A simpler Arabidopsis protoplast isolation method. Plant Methods, 5(1), 16. 10.1186/1746-4811-5-16

Xu, L., Wu, C., Oelmüller, R., & Zhang, W. (2018). Role of Phytohormones in Piriformospora indica-Induced Growth Promotion and Stress Tolerance in Plants: More Questions Than Answers. Frontiers in Microbiology, 9. https://www.frontiersin.org/articles/10.3389/fmicb.2018.01646

Yoo, S.-D., Cho, Y.-H., & Sheen, J. (2007). Arabidopsis mesophyll protoplasts: A versatile cell system for transient gene expression analysis. Nature Protocols, 2(7), 1565–1572. 10.1038/nprot.2007.199

Zand Karimi, H., & Innes, R. W. (2022). Molecular mechanisms underlying host-induced gene silencing. The Plant Cell, 34(9), 3183–3199. 10.1093/plcell/koac165

Zhan, J., & Meyers, B. C. (2023). Plant Small RNAs: Their Biogenesis, Regulatory Roles, and Functions. Annual Review of Plant Biology, 74(1), 21–51. 10.1146/annurev-arplant-070122-035226

Zhang, T., Zhao, Y.-L., Zhao, J.-H., Wang, S., Jin, Y., Chen, Z.-Q., Fang, Y.-Y., Hua, C.-L., Ding, S.-W., & Guo, H.-S. (2016). Cotton plants export microRNAs to inhibit virulence gene expression in a fungal pathogen. Nature Plants, 2(10), 16153. 10.1038/nplants.2016.153

Zuccaro, A., Lahrmann, U., Güldener, U., Langen, G., Pfiffi, S., Biedenkopf, D., Wong, P., Samans, B., Grimm, C., Basiewicz, M., Murat, C., Martin, F., & Kogel, K.-H. (2011). Endophytic Life Strategies Decoded by Genome and Transcriptome Analyses of the Mutualistic Root Symbiont Piriformospora indica. PLOS Pathogens, 7(10), e1002290. 10.1371/journal.ppat.1002290

