## Supplemental File 1 for "A pipeline for validation of sRNA effectors suggests cross-kingdom communication in the symbiosis of *Arabidopsis* with *Serendipita indica*"

**Supplemental Table 1:** Sequences of amiRNAs cloned in pUC18-*At*MIR390a-B/c construct and expressed in *At* protoplasts for PTGS validation, their 5´- nucleotides, their corresponding predicted Arabidopsis target genes, and their target predicted scores. Artificial microRNAs (amiRNAs) are designed using P.SAMS software (https://p-sams.carringtonlab.org) developed by the James Carrington lab. An expectation value indicates the penalty of mismatches between amiRNAs and the target sequence (*At* predicted target) with an expectation value of 0 being the highest level of sequence complementarity and 5 being the lowest.

| sRNA name | sRNA sequence | 5´-terminal nucleotide | sRNA corresponding target | Target aligned fragment | Expectation value |
|---|---|---|---|---|---|
| amir21 | UACAUUCGCUCAGGUUCACCA | U | *AT5G25350* | AGGUGAACCUGAGCGAAUGUA | 0 |
| amir24 | UCAAUCUCUAGAUGUUGACCA | U | *AT2G39020* | AGGUCAACAUCUAGAGAUUGA | 0 |
| amir154 | UCCCUAGGAUAUUAUUGACAA | U | *AT4G32160* | AUGUCAAUAAUAUCCUAGGGA | 0 |
| amir296 | UUGUUCAGGCGUUUUUAUCU | U | *AT2G45240* | AGAUAAAAACGCCUGAACAA | 0 |

**Supplemental Table 2:** Sequences of *Si*sRNA cloned in pUC18-*At*MIR390a-B/c construct and expressed into *At* protoplasts for PTGS validation, their 5´- nucleotides, their corresponding predicted Arabidopsis target genes, and their expectation values indicating the degree of mismatches with the predicted *At* target sequence as provided by psRNAtarget (Dai et al., 2018). An expectation value indicates the penalty of mismatches between mature small RNA (*Si*sRNA) and the target sequence (*At* predicted target) with a value of 0 being the highest level of sequence complementarity and 5 being the lowest.

| sRNA name | sRNA sequence | 5´-terminal nucleotide | predicted Arabidopsis target gene | Target aligned fragment | Expectation value |
|---|---|---|---|---|---|
| *Si*sRNA21 | CAAAUUUUGAAUCUGGCGCCU | C | *AT1G01210* | AUGAGCCAGAAUCAAGAUUUU | 4 |
| | | | *AT5G25350* | AGGCGACGGAUUUGAGGUUGG | 5 |
| | | | *AT5G37500* | GAGUGUCAGAUUCAAGAGUUG | 3 |

| | | | | | |
|---|---|---|---|---|---|
| | | | *AT5G37600* | UCUUGCAAGAUUCAAAGUUUG | 3 |
| *Si*sRNA24 | UUGACCAAUUUUUCUGUUCUC | U | *AT1G57590* | GAGGAAAGAGAAUUUGGUUAG | 4.5 |
| | | | *AT1G63180* | GAGAGUGGAAGAAUUGGAGAA | 5 |
| | | | *AT1G65090* | GAGUAUAGCAAGAUUGGUCAC | 4.5 |
| | | | *AT4G15765* | AAGAAUGGAAGAAUUGGACAU | 4 |
| | | | *AT5G16680* | GAGAACAGGAAGGAUGGUAAG | 5 |
| *Si*sRNA28 | ACAACUUUCAACAACGGAUCU | A | *AT1G05180* | AGAUUCAAUGUUGAAACUUGA | 5 |
| | | | *AT3G06670* | CGAUUUGGUGUUGGAGGUUGG | 4 |
| | | | *AT5G55930* | AGGUGUUGUUGUUGAAGGUUGU | 4 |
| *Si*sRNA154 | UCUACCUCUUGGUGAGCCGGC | U | *AT2G47600* | GUUGACUUAUUAGGAGGUGGG | 4.5 |
| | | | *AT4G32160* | GUCUGCAUGCCAAGAUGUAGA | 4.5 |

**Supplemental Table 3:** 75 mer oligonucleotides used for cloning *Si*sRNAs and amiRNAs into the pUC18-AtMIR390a-RFP vector.

| Primer name | 75mer sequence |
|---|---|
| amir296-*AT2G45240*-75-F | TGTATTGTTCAGGCGTTTTTATCTGATGATGATCACATTCGTTATCTATTTTTTCAGATAAAAAGCCTGAACAA |
| amir296-*AT2G45240*-75-R | AATGTTGTTCAGGCTTTTTATCTGAAAAAATAGATAACGAATGTGATCATCATCAGATAAAAACGCCTGAACAA |
| amir24-*AT2G39020*-75-F | TGTATCTTCACCGCTTGTTTGGCCTATGATGATCACATTCGTTATCTATTTTTAGGCCAAACACGCGGTGAAGA |
| amir24-*AT2G39020*-75-R | AATGTCTTCACCGCGTGTTTGGCCTAAAAAATAGATAACGAATGTGATCATCATAGGCCAAACAAGCGGTGAAGA |
| amir21-*AT5G25350*-75-F | TGTATACATTCGCTCAGGTTCACCAATGATGATCACATTCGTTATCTATTTTTTGGTGAACCTTAGCGAATGTA |
| amir21 -*AT5G25350* -75 -R | AATGTACATTCGCTAAGGTTCACCAAAAAAATAGATAACGAATGTGATCATCATTGGTGAACCTGAGCGAATGTA |
| amir154 -*AT4G32160* -75 -F | TGTATCCCTAGGATATTATTGACAAATGATGATCACATTCGTTATCTATTTTTTTTGTCAATAAGATCCTAGGGA |
| amir154 -*AT4G32160* -75 -R | AATGTCCCTAGGATCTTATTGACAAAAAAAATAGATAACGAATGTGATCATCATTTGTCAATAATATCCTAGGGA |
| *Si*sRNA21 -75 -F | TGTACAAATTTTGAATCTGGCGCCTATGATGATCACATTCGTTATCTATTTTTAGGCGCCAGAGTCAAAATTTG |
| *Si*sRNA21 -75 -R | AATGCAAATTTTGACTCTGGCGCCTAAAAAATAGATAACGAATGTGATCATCATAGGCGCCAGATTCAAAATTTG |
| *Si*sRNA24 -75 -F | TGTATTGACCAATTTTTCTGTTCTCATGATGATCACATTCGTTATCTATTTTTTGAGAACAGAACAATTGGTCAA |

| | |
|---|---|
| *Si*sRNA24 -75 -R | AATGTTGACCAATTGTTCTGTTCTCAAAAAATAGATAACGAATGT GATCATCATGAGAACAGAAAAATTGGTCAA |
| *Si*sRNA28 -75 -F | TGTAACAACTTTCAACAACGGATCTATGATGATCACATTCGTTAT CTATTTTTTAGATCCGTTGGTGAAAGTTGT |
| *Si*sRNA28 -75 -R | AATGACAACTTTCACCAACGGATCTAAAAAATAGATAACGAATG TGATCATCATAGATCCGTTGTTGAAAGTTGT |
| *Si*sRNA154 -75 -F | TGTATCTACCTCTTGGTGAGCCGGCATGATGATCACATTCGTTA TCTATTTTTTGCCGGCTCACAAAGAGGTAGA |
| *Si*sRNA154 -75 -R | AATGTCTACCTCTTTGTGAGCCGGCAAAAAATAGATAACGAATG TGATCATCATGCCGGCTCACCAAGAGGTAGA |

**Supplemental Table 4:** Primers used in Stem-loop PCR (cDNAhp = complementary DNA hairpin, F = forward)

| Primer name | Primer sequence |
|---|---|
| amir296-*AT2G45240*-cDNAhp | GTCGTATCCAGTGCAGGGTCCGAGGTATTCGCACTGGATACGACc agata |
| amir296-*AT2G45240*-F | TCGCTttgttcaggcgtttt |
| amir24-*AT2G39020*-cDNAhp | GTCGTATCCAGTGCAGGGTCCGAGGTATTCGCACTGGATACGACa ggcca |
| amir24-*AT2G39020*-F | TCGCTtcttcaccgcttgtt |
| amir21-*AT5G25350*-cDNAhp | GTCGTATCCAGTGCAGGGTCCGAGGTATTCGCACTGGATACGACt ggtga |
| amir21-*AT5G25350*-F | TCGCTtacattcgctcaggt |
| amir154-*AT4G32160*-cDNAhp | GTCGTATCCAGTGCAGGGTCCGAGGTATTCGCACTGGATACGACt tgtca |
| amir154-*AT4G32160*-F | TCGCTtccctaggatattat |
| *Si*sRNA21-cDNAhp | GTCGTATCCAGTGCAGGGTCCGAGGTATTCGCACTGGATACGACa ggcgc |
| *Si*sRNA21-F | TCGCTcaaattttgaatctg |
| *Si*sRNA24-cDNAhp | GTCGTATCCAGTGCAGGGTCCGAGGTATTCGCACTGGATACGACg agaac |
| *Si*sRNA24-F | TCGCTttgaccaatttttct |
| *Si*sRNA28-cDNAhp | GTCGTATCCAGTGCAGGGTCCGAGGTATTCGCACTGGATACGACa gatcc |
| *Si*sRNA28-F | TCGCTacaactttcaacaac |
| *Si*sRNA-154-cDNAhp | GTCGTATCCAGTGCAGGGTCCGAGGTATTCGCACTGGATACGACg ccggc |
| *Si*sRNA154-F | TCGCTtctacctcttggtga |
| *At*miR159a-cDNAhp | GTCGTATCCAGTGCAGGGTCCGAGGTATTCGCACTGGATACGACt agagc |
| *At*miR159a-F | TCGCTtttggattgaaggga |
| Univ-stemloop-reverse | GTATCCAGTGCAGGGTCCGAGGT |

**Supplemental Table 5:** Primers used for gene expression analysis in quantitative real-time PCR (qPCR).

| Primer name | Primer sequence |
|---|---|

| | |
|---|---|
| q*AT1G01210* -F | CTGAAACTGAAGCCCCATGT |
| q*AT1G01210* -R | ATTCCTCACGCCAAGTGAAC |
| q*AT5G25350* -F | AGGAGTTTGGAGGGGAAGAA |
| q*AT5G25350* -R | ATGGACAACCATGAGCAACA |
| q*AT5G37600* -F | CAATCCTCTGGAATCCTTGA |
| q*AT5G37600* -R | AAAAGCAGAATAAGCAGAGCAAA |
| q*AT1G57590* -F | TTGTGACGGTGGATCGTTTA |
| q*AT1G57590* -R | AGCAGAGCCTGCTTAGCTTG |
| q*AT1G63180* -F | GCGGAACCAGAATGGAAGAT |
| q*AT1G63180* -R | TTATTCGGTATGCCCTTTGG |
| q*AT1G65090* -F | AGTTGGCAAGGAGGATGATG |
| q*AT1G65090* -R | TGAATGGGAATGGGTTCTTC |
| q*AT4G15765* -F | CGACATTGCAGGACATAACG |
| q*AT4G15765* -R | CCTGTGAAGAGCAAGGGAAG |
| q*AT5G16680* -F | TTTGCAGAATCCATCATCCA |
| q*AT5G16680* -R | TCCTGAACGCGTAGATCCTT |
| q*AT2G39020* -F | TTTGTCCAAGCCTGGTTTTT |
| q*AT2G39020* -R | TTCCAATCAAGAACAACCCATT |
| qA*T1G05180* -F | AGTGTTGGCCAATCAAAAGC |
| q*AT1G05180* -R | GCCATAAGAGCGAACCAAAA |
| q*AT3G06670* -F | CTCGTTGCTCCAAAACCCTA |
| q*AT3G06670* -R | GGAGCGCCCATAACTGATTA |
| q*AT5G55930* -F | GGGCTTGAGAATTCAGACCA |
| q*AT5G55930* -R | TTACCCCTTTCAGGACAACG |
| q*AT2G47600* -F | CTCCGATTCTTTTCCCCAGT |
| q*AT2G47600* -R | TTCTTGCAACTCCTGGGTTT |
| q*AT4G32160* -F | GGTGCTCTCTTGAGGAGTGG |
| q*AT4G32160* -R | ATAGGCGACGAACTTGTGCT |
| q*AT2G45240* -F | AAGGCCAGGCTAGAACACCT |
| qA*T2G45240* -R | TCACGCATTCTCTGGATTTG |
| qUbiquitin -F | CCAAGCCGAAGATCAAG |
| qUbiquitin -R | ATGACTCGCCATGAAAGTCC |
| qActin -F | GGAAGGATCTGTACGGTAAC |
| qActin -R | TGTGAACGATTCCTGGACCT |

**Supplemental Table 6:** Primers used for RNA ligase-mediated rapid amplification of cDNA ends (RLM-RACE)

| Primer Name | Primer sequence |
|---|---|
| RLM -*AT2G45240* -Outer -Specific | ATCAGGCCATGTTCGATCCC |
| RLM -*AT2G45240* -Inner -Specific | TTGTAGCATGGCGGTTGAC |
| RLM -*AT1G15765* -Outer -Specific | TTAACGTGCACAATCGGAAA |
| RLM -*AT1G15765* -Inner -Specific | GCCAAGAGGAAGAGCATGAG |
| RLM -*AT1G65090* -Outer -Specific | CCTCATCGTCCTTGCCCGTAGAA |
| RLM -*AT1G65090* -Inner -Specific | CTTTACCCCTGAAAAGTGCAGCC |
| RLM -Outer -Universal | GCTGATGGCGATGAATGAACACTG |
| RLM -Inner -Universal | GAACACTGCGTTTGCTGGCTTTGATG |

**Supplemental Table 7:** PCR primers

| Primer Name | Primer sequence |
|---|---|
| ITS -Fwd | CAACACATGTGCACGTCGAT |
| ITS -Rev | CCAATGTGCATTCAGAACGA |
| *Si* -Ubi -Fwd | GCAGCTCGAAGATGGTCGC |
| *Si* -Ubi -Rev | ACATGCACGCTTGCGGAGT |
| M13 -Fwd | GTTTTCCCAGTCACGAC |
| M13 -Rev | AACAGCTATGACCATG |
| Attb -Fwd | GGGGACAAGTTTGTACAAAAAAGCAGGCT |
| Attb -Rev | GGGGACCACTTTGTACAAGAAAGCTGGGT |
| pUC18-Mut-Fwd | GCAATGATACCGCGAGAGCCACGCTCACCGGCTCC |
| pUC18-Mut-Rev | GGAGCCGGTGAGCGTGGCTCTCGCGGTATCATTGC |

Red nucleotides present the substituted bases for site-directed mutagenesis incorporated in the restriction site of BsaI in the backbone of the pUC18 vector (see material and methods).

**Supplementary Table 8:** Sequences, 5´-terminal nucleotide, raw read counts normalized to *Si*-colonized *Bd* roots or to *Si* axenic, and the induction status of the 14 selected putative ck-*Si*sRNAs from our previous study (Šečić *et al.,* 2021) investigating *Si*sRNA expression profiles during the mutualistic interaction between *Serendipita indica* and the grass model *Brachypodium distachyon* using high throughput sRNAseq.

| ***Si*sRNA name** | ***Si*sRNA sequence** | **5´- terminal nucleotide** | **Raw reads** | **Raw reads normalized to *Si-Bd* colonized** | **Raw reads normalized to *Si* axenic** | **Induction** |
|---|---|---|---|---|---|---|
| ***Si*sRNA21** | **CAAATTTTGAATCTGGCGCCT** | **C** | **212** | **165,27** | **6,01** | **up** |
| ***Si*sRNA23** | **TACCCATACCTCGCCGTCGGC** | **T** | **95** | **74,06** | **4,42** | **up** |
| ***Si*sRNA24** | **TTGACCAATTTTTCTGTTCTC** | **T** | **57** | **44,43** | **4,56** | **up** |
| *Si*sRNA25 | TTGGAAGCGGCTGGACTAGTC | T | 11 | 8,57 | 1,00 | up |
| *Si*sRNA26 | TACAACTTTCAACAACGGATC | T | 82 | 63,92 | 14,73 | up |
| ***Si*sRNA28** | **ACAACTTTCAACAACGGATCT** | **A** | **595** | **463,85** | **109,6** | **up** |
| *Si*sRNA37 | GCTCACGTTCTATAGATTTGT | G | 330 | 257,26 | 70,28 | up |
| *Si*sRNA47 | GGCCATGGAAGTCGGAACCCG | G | 6 | 4,67 | 1,23 | up |
| *Si*sRNA62 | ATCCACGGCCATAGGACTCTG | A | 2608 | 2033,18 | 856,15 | up |
| *Si*sRNA86 | CCGGGGTGTATTTATTAGATA | C | 3 | 2,33 | 1,26 | up |
| ***Si*sRNA154** | **TCTACCTCTTGGTGAGCCGGC** | **T** | **35** | **27,28** | **0.47** | **up** |
| *Si*sRNA165 | TCGGGCAATGGCGCGAAGCTA | T | 7 | 5,45 | 0.82 | up |
| ***Si*sRNA296** | **TCTGCGAATCGTAATTAAATA** | **T** | **2** | **1,55** | **0.26** | **up** |
| *Si*sRNA376 | TGATGCTCAGAACGGCGGCTC | T | 2 | 1,55 | #N/A | exclusive |

*Si*sRNAs confirmed by Stem-Loop PCR and Sanger sequencing to be expressed in 4-week-old axenic culture are highlighted **in bold**.

*Si*sRNAs confirmed by Stem-Loop PCR and Sanger sequencing to be expressed in *Si*-colonized *At* roots are Highlighted **in bold and blue**.

**Supplemental Table 9:** The expression level of *Arabidopsis thaliana* target genes used in our study in transcript per million (TPM) in leaf and root. Data extracted from (Moreno et al., 2022)

| Gene ID | Description | Expression level in leaf | Expression level in root |
|---|---|---|---|
| *AT2G39020* | Acyl-CoA N-acyltransferases (NAT) superfamily protein \| Chr2:16295140-16296495 FORWARD LENGTH=1356 \| 201606 | 40.0 | 48.0 |
| *AT2G45240* | methionine aminopeptidase 1A \| Chr2:18655760-18659106 FORWARD LENGTH=1696 \| 201606 | 27.0 | 52.0 |
| *AT1G01210* | DNA-directed RNA polymerase, subunit M, archaeal \| chr1:88898-89745 FORWARD LENGTH=616 | 10.0 | 16.0 |
| *AT5G25350* | EIN3-binding F box protein 2 \| chr5:8794252- | 53.0 | 119.0 |

| | | | |
|---|---|---|---|
| | 8797020 REVERSE LENGTH=2600 | | |
| *AT5G37500* | gated outwardly-rectifying K+ channel \| chr5:14889692-14895214 REVERSE LENGTH=2860 | 4.0 | 13.0 |
| *AT5G37600* | glutamine synthase clone R1 \| chr5:14933336-14935841 REVERSE LENGTH=1494 | 35.0 | 293.0 |
| *AT1G57590* | Pectinacetylesterase family protein \| chr1:21327360-21329763 REVERSE LENGTH=1489 | 1.0 | 24.0 |
| *AT1G63180* | UDP-D-glucose/UDP-D-galactose 4-epimerase 3 \| chr1:23427409-23429603 REVERSE LENGTH=1425 | 20.0 | 37.0 |
| *AT1G65090* | unknown protein; FUNCTIONS IN: molecular_function unknown; INVOLVED IN: biological_process unknown; LOCATED IN: cellular_component unknown; BEST Arabidopsis thaliana | NA | 0.4 |

| | | | |
|---|---|---|---|
| | protein match is: unknown protein (TAIR:AT5G36100.1). \| chr1:24180876-24183300 REVERSE LENGTH=1200 | | |
| *AT4G15765* | FAD/NAD(P)-binding oxidoreductase family protein \| Chr4:8975281-8977434 REVERSE LENGTH=1658 \| 201606 | 4.0 | 8.0 |
| *AT5G16680* | RING/FYVE/PHD zinc finger superfamily protein \| Chr5:5467331-5473054 REVERSE LENGTH=4237 \| 201606 | 21.0 | 40.0 |
| *AT1G05180* | NAD(P)-binding Rossmann-fold superfamily protein \| chr1:1498114-1501824 REVERSE LENGTH=1817 | 24.0 | 48.0 |
| *AT3G06670* | binding \| chr3:2105459-2113421 REVERSE LENGTH=3332 | 42.0 | 53.0 |
| *AT5G55930* | oligopeptide transporter 1 \| chr5:22652715- | 12.0 | 0.3 |

| | | | |
|---|---|---|---|
| | 22656106 FORWARD LENGTH=2820 | | |
| *AT2G47600* | magnesium/proton exchanger \| Chr2:19524117-19527580 REVERSE LENGTH=2357 \| 201606 | 20.0 | 37.0 |
| *AT4G32160* | Phox (PX) domain-containing protein \| Chr4:15528425-15533279 FORWARD LENGTH=3190 \| 201606 | 16.0 | 38.0 |

**Supplemental Table 10:** Sequences of *Si*sRNA detected in *Si* axenic and *Si-At* interaction compared to the respective stem-loop PCR amplicons detected from *Si-At* Co-IP.

| sRNA name | ***Si*sRNA sequence detected in *Si* axenic and/or *Si-At*** | **Sequence of *Si*sRNA detected in Co-IP** |
|---|---|---|
| *Si*sRNA21 | CAAATTTTGAATCTGG CGCCT | TCGCT**CAAATTTTGAATCTGTCGATC**CAGTGCGAATACCTC GGACCCTGCACTGGATACAATCGAATTCCCGCGGCCGCC |
| *Si*sRNA23 | TACCCATACCTCGCCG TCGGC | TCGCT**TACCCATACCTCGCCGTCGTA**TCCAGTGCGAATACCT CGGACCCTGCACTGGATACAATCACTAGTGAATTCGCG |
| *Si*sRNA24 | TTGACCAATTTTTCTGT TCTC | TCGCT**TTGACCAATTTTTCTGTTCTC**GTCCGTATCCAGTGCG AATACCTCGGACCCTGCACTGGATACAATCGAATTCCC |
| *Si*sRNA28 | ACAACTTTCAACAACG GATCT | TCGCT**ACAACTTTCAACAACGGATCT**GTCGTATCCAGTGCG AATACCTCGGACCCTGCACTGGATACAATCACTAGTGAA |
| *At*miR159a | TTTGGATTGAAGGGA GCTCTA | GCGCC**TTTGGATTGAAGGGAGCTCTA**GTCGTATCCAGTGC GAATACCTCGGACCCTGCACTGGATACAATCGAATTCCCG |

**Bold letters** highlight the presence of the 21 nt *Si*sRNA sequences (detected in *Si* axenic and *Si-At*) in the stem-loop PCR amplicon detected from Co-IP. **Red letters** indicate mismatches of the Co-IP sequencing results relative to the sequencing results of *Si*sRNA in the axenic culture.

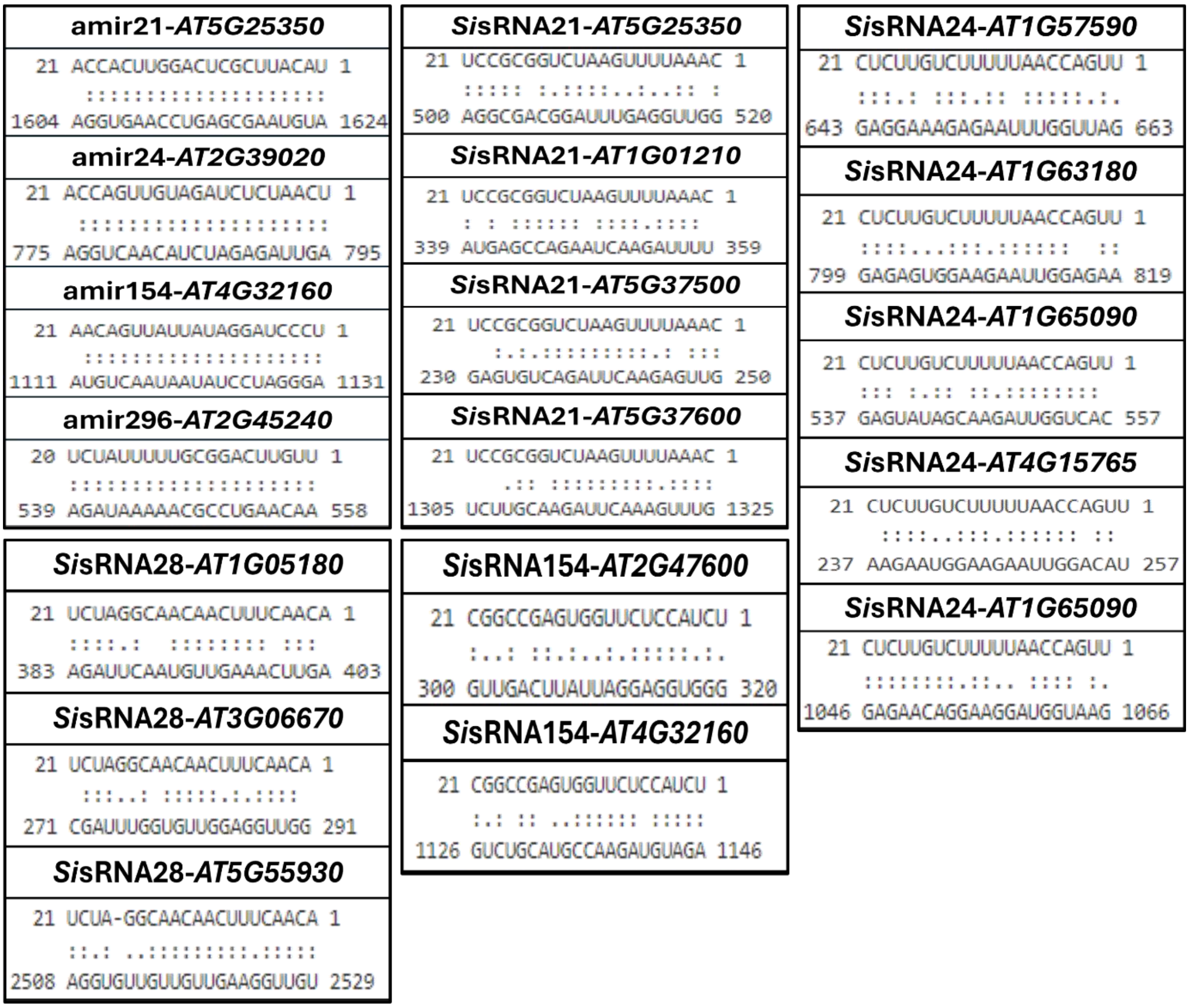


**Supplemental Figure 1:** Illustration of the alignment between the investigated amiRNAs and *Sis*RNA and their predicted Arabidopsis target genes as displayed by psRNATarget web tool.

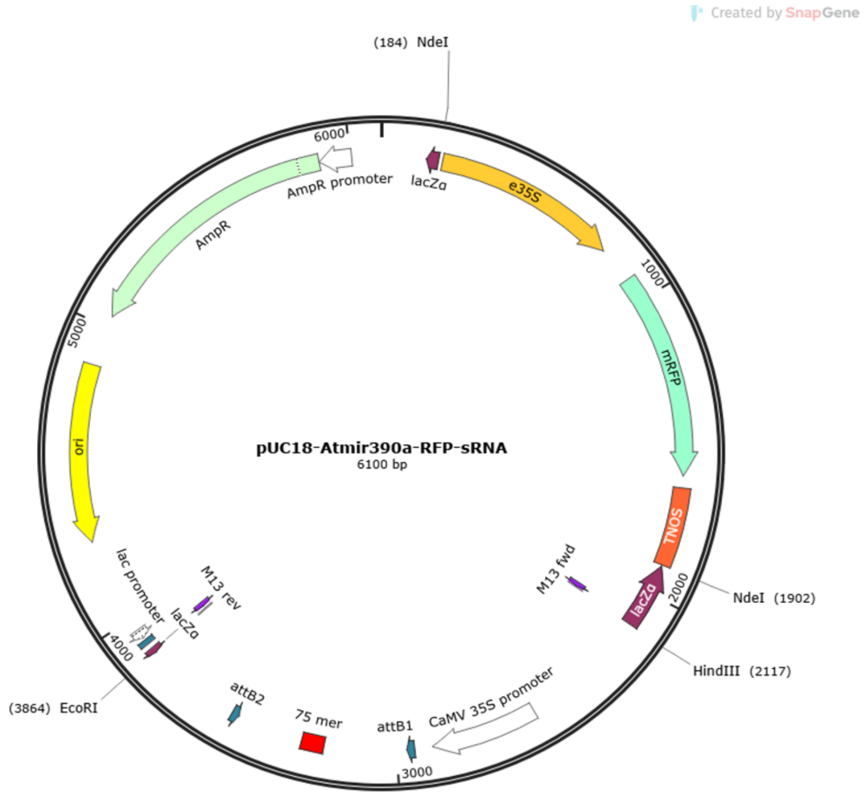


**Supplemental Figure 2. *In-silico* cloning of a pUC18-based vector for the expression of amiRNAs and *Sis*RNAs.** An *At*MIR390a-B/c (*At*MIR390a-*Bsa*I/*ccdB*) fragment was subcloned from the pMDC32B-*At*MIR390a-B/c vector (Plasmid #51776, Carbonell et al., 2014) into pUC18 (Plasmid #50004), using *Eco*RI and *Hind*III, resulting in pUC18 -*At*MIR390a-B/c. The *CaMV35S::RFP* expression cassette was cloned from the pBeaconRFP vector (https://gatewayvectors.vib.be/index.php/ ID: 3_20, Bargmann and Birnbaum Plant Physiol. (2009)) into pUC18-*At*MIR390a-B/c using *Nde*I, resulting in pUC18-*At*MIR390a-B/c-RFP. The *Bsa*I restriction site in the backbone of pUC18-*At*MIR390a-B/c-RFP was removed by site-directed mutagenesis. 75mer oligos encoding 21nt sRNA sequences (Suppl. Table 1 & 2) were annealed and cloned into the *Bsa*I sites of pUC18-*At*MIR390a-B/c-RFP by GoldenGate cloning, resulting in the final pUC18-*At*MIR390a-RFP-sRNA expression vector. *In-silico* cloning was created with SnapGene software. Vector maps are available at https://www.addgene.org/.

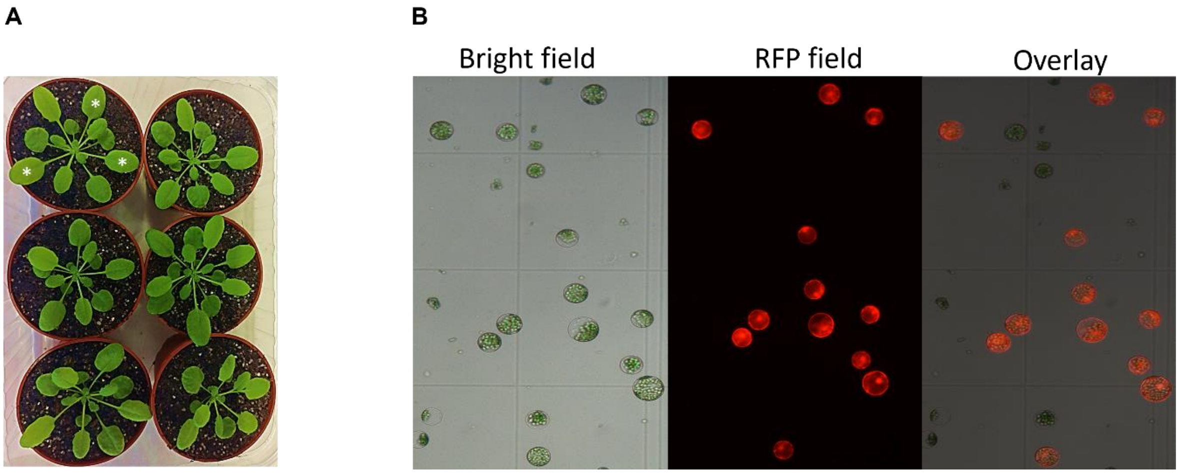


**Supplementary figure 3. Expression of amiRNAs or *Si*sRNAs in Arabidopsis leaf protoplasts (A)** 4- to 5-week-old Arabidopsis (Col-0) plants suitable for protoplast isolation. (*) optimal leaves to be used for protoplast isolation. **(B)** Arabidopsis leaf protoplasts 1 day after transformation with the expression vector pUC18-*At*MIR390-RFP-sRNA containing a red fluorescing protein (RFP) gene under the control of the CaMV35S promoter to confirm successful transformation. Red protoplasts in the RFP image carry the expression vector.

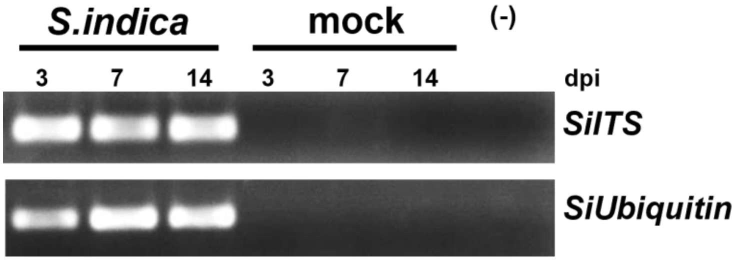


**Supplemental Figure 4. Expression of *S. indica*-specific gene and the internal transcribed spacer (ITS) in Arabidopsis roots inoculated with *S. indica* as analyzed by semi-quantitative reverse-transcription (RT) PCR. (A)** Expression of the Internal Transcribed Spacer (Si*ITS*) and **(B)** of *SiUbiquitin* at 3-, 7- and 14 days post inoculation (dpi) of Arabidopsis roots with *Si* chlamydospores. Arabidopsis roots treated with $H_2O_{0.02\%\ Tween\ 20}$ were used as a control (mock). (-) = non template control

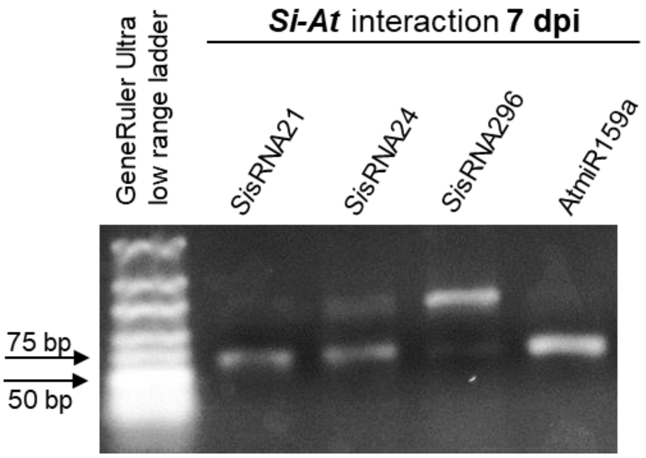


**Supplemental Figure 5. Detection of *Si*sRNAs by stem-loop PCR in Arabidopsis roots 7 days after inoculation (dpi) with *S. indica*.** *At*miR159a was used as an endogenous control.

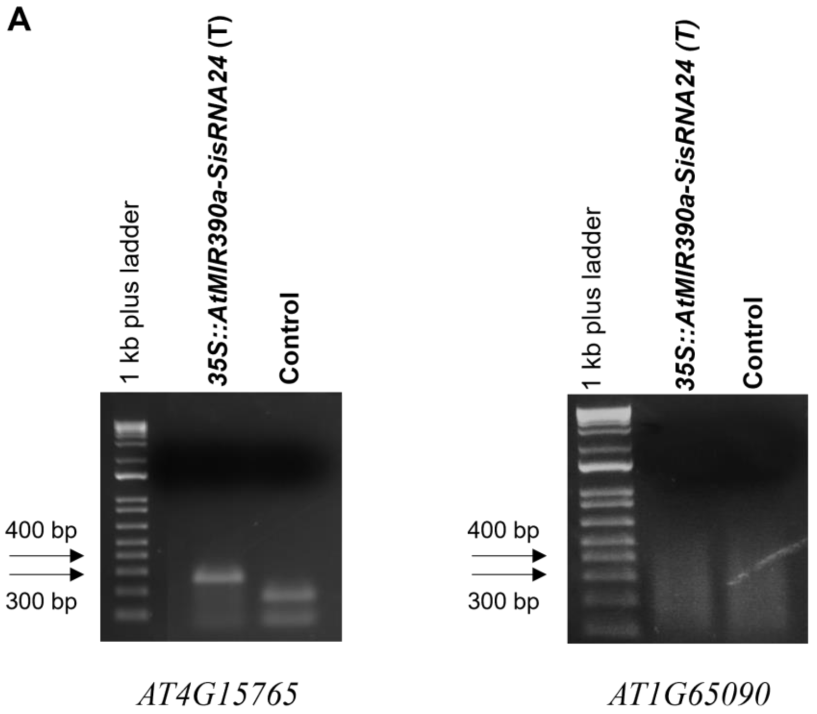

**B**

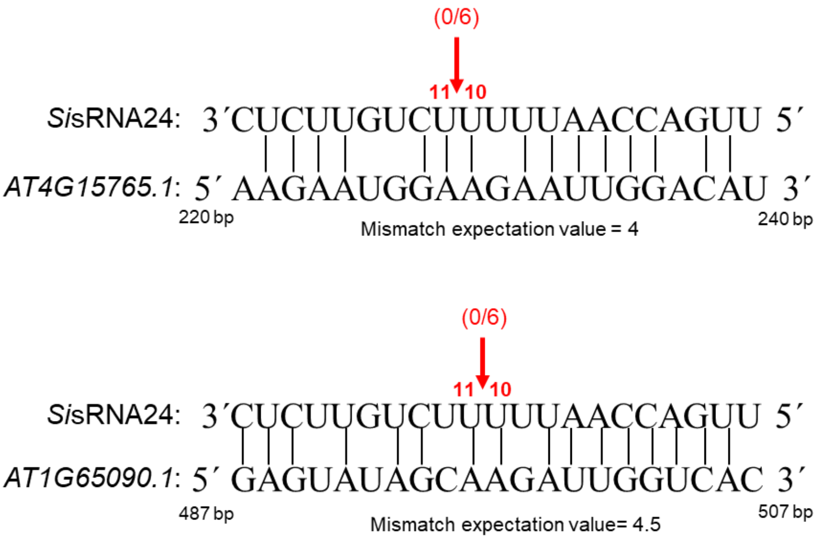


**Supplemental figure 6. 5′ RLM-RACE for two Arabidopsis target genes of *Si*sRNA24 after expression in Arabidopsis protoplasts.** RNA was extracted from Arabidopsis protoplasts 24 h after transformation with an expression construct of *Si*sRNA24. Water replaced *Si*sRNA24 in protoplasts and served as control. **(A)** PCR amplification after 5′ RLM-RACE for the two *Si*sRNA24 targets *AT4G15765* and *AT1G65090* as visualized by Agarose gel electrophoresis. No clear PCR amplicon was detected for *AT1G65090* **(B)** Predicted mapping of *AT4G15765* and *AT1G65090* cleavage sites. Red arrows indicate the expected canonical cleavage sites. The proportion of cloned 5´-RLM-RACE products at the predicted cleavage sites is shown in brackets. Expectation value is provided by the psRNATarget server (http://plantgrn.noble.org/psRNATarget/). An expectation value indicates the penalty of mismatches between mature small RNA (*Si*sRNA) and the target sequence (*At* predicted target) with an expectation value of 0 being the highest level of sequence complementarity and 5 being the lowest.

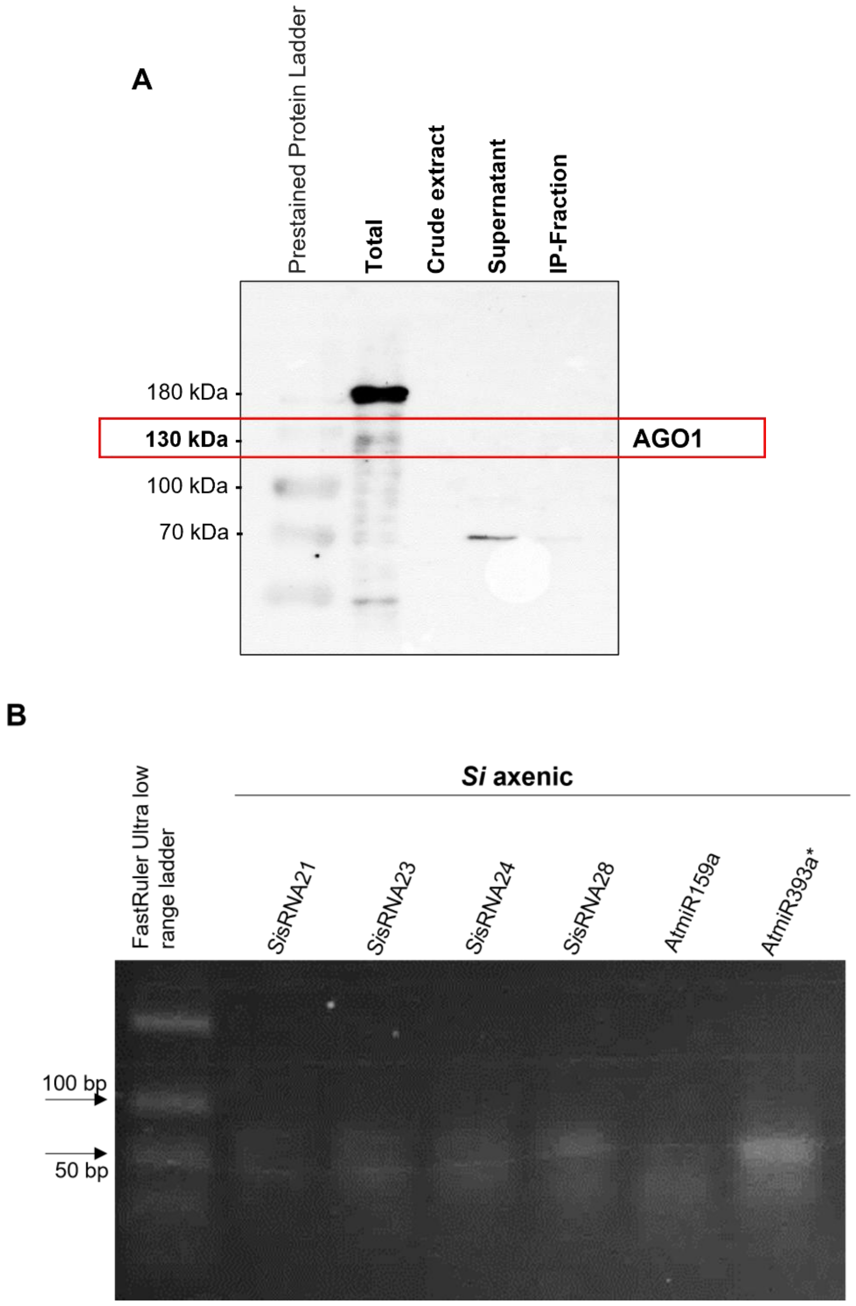


**Supplementary figure 7. *At*AGO1/sRNA co-immunoprecipitation from *Si* axenic culture to show the specificity of the Arabidopsis AGO1 antibody. (A)** Total protein was isolated from *S. indica* grown in axenic culture for 4 weeks and immunoprecipitation was performed using an anti-*At*AGO1 antibody. No signal could be detected in the *Si* samples excluding cross-reaction of the anti-*At*AGO1 antibody with *Si* AGO proteins. Weak unspecific bands of 70 kDa were detected in the SN and the IP fraction suggesting an unspecific binding to unrelated proteins or the presence of a contaminant. **(B)** Stem-loop PCR to detect *Si*sRNAs recovered from *At*AGO1/sRNA co-immunoprecipitation from *Si* axenic culture. No amplification or weak bands of primer dimers were obtained. Arabidopsis *At*miR159a and *At*miR393a* were used as negative controls.

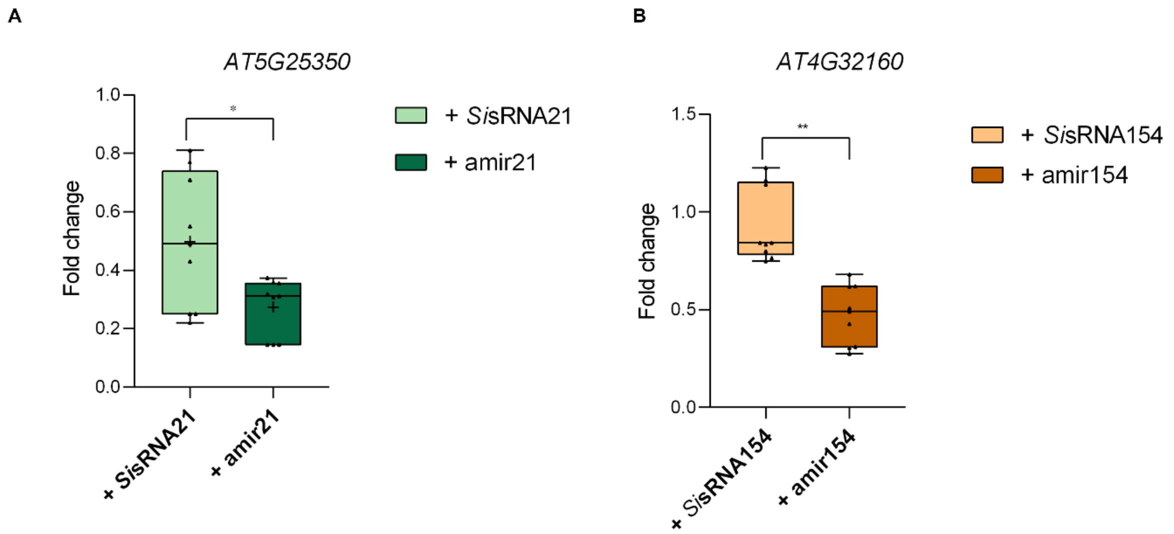


**Supplemental Figure 8. Comparison of putative and artificial sRNA and their mediated down-regulation after their transient expression in Arabidopsis protoplasts. (A)** comparison of *Si*sRNA21 and amir21 and **(B)** of *Si*sRNA154 and amir154. Arabidopsis protoplasts were transformed with either pUC18-*At*MIR390a-RFP-*Si*sRNA or pUC18-*At*MIR390a-RFP-amiRNA. Control protoplasts were treated with water. Total RNA was extracted 24 h after transformation and reverse transcribed into cDNA. The Arabidopsis housekeeping gene *UBIQUITIN* (*AT3G62250*) was used for normalization. Data show fold changes in the expression of *AT5G25350* and *AT4G32160* relative to the control sample as analyzed by quantitative Real-Time PCR. RT-PCR. Bars represent the average of three independent biological replicates ± standard deviation. The asterisks indicate a significant difference at $P < 0.05$ according to an unpaired Student's *t-test*.
